# A one-track model for spatiotemporal coordination of *Bacillus subtilis* septal cell wall synthesis

**DOI:** 10.1101/2023.06.29.547024

**Authors:** Kevin D. Whitley, James Grimshaw, David M. Roberts, Eleni Karinou, Phillip J. Stansfeld, Séamus Holden

## Abstract

Bacterial cell division requires synthesis of a septal peptidoglycan (sPG) wall across the middle of the cell. This is accomplished by the divisome synthesis complex in coordination with numerous other division proteins—such as the essential tubulin homolog FtsZ—but the molecular mechanism of its spatiotemporal regulation remains unclear. Here, we investigate the dynamics of sPG synthesis in the model Gram-positive bacterium *Bacillus subtilis* using live-cell single-molecule imaging of the divisome transpeptidase PBP2B. In contrast to previous models for division, we show that there is a single population of processively-moving PBP2B molecules whose motion is driven by peptidoglycan synthesis and is not associated with FtsZ treadmilling. However, although the motions of PBP2B and FtsZ are asynchronous, we demonstrate that processive PBP2B motion is partially dependent on FtsZ treadmilling. Additionally, we provide evidence that the divisome synthesis complex is multimeric. Our results support a new model for division in *B. subtilis* where a multimeric synthesis complex follows a single track dependent on sPG synthesis whose activity and dynamics are asynchronous with FtsZ treadmilling.

## Introduction

Cell division is a basic requirement for bacterial life and a major antibiotic target. At a molecular level, division is a remarkable feat of engineering by the bacterial cell: a set of nanoscale proteins must cooperate over large distances to build a micron-scale cross-wall (septum) across the middle of the cell against heavy outward pressure. Many of the proteins involved are highly conserved across the bacterial domain, including the septal peptidoglycan (sPG) synthases that insert new cell wall material into the ingrowing septum and the cytoskeletal tubulin homolog FtsZ^1^. FtsZ forms short filaments that move around the division septum by treadmilling^2–5^ – a type of motion where monomers bind one end of a filament at the same rate they unbind the opposite end, causing the filament to move forward even though monomers remain stationary. In contrast, sPG synthases are part of a larger divisome synthesis complex^6^ that does not treadmill, but instead moves processively around the division ring^2,3,5,7^.

Several models have been proposed to explain how these proteins cooperate to enact division despite such different motion patterns^8^. We previously proposed a model where FtsZ treadmilling drives septal constriction in *B. subtilis* as a coupled cytoskeleton-synthase complex^2^. However, subsequent work in this and other Bacillota (a.k.a. Firmicute) species demonstrated that sPG synthesis is not tightly coupled to FtsZ treadmilling. FtsZ treadmilling is dispensable for septal constriction after constriction has initiated in both *Staphylococcus aureus* and *B. subtilis*^4,9^, and the motions of divisome synthesis complexes are uncoupled from treadmilling FtsZ filaments in *Streptococcus pneumoniae*^5^. Meanwhile, work on the Pseudomonadota (a.k.a. Proteobacteria) species *Escherichia coli* and *Caulobacter crescentus* has supported a model where the synthase complex moves on two ‘tracks’: an FtsZ-track where inactive synthase complexes are distributed around the division septum and an sPG-track where synthase complexes build the cell wall independently of FtsZ^10–13^. According to this model, an activating protein (FtsN in *E. coli* and FzlA in *C. crescentus*) is required to initiate active sPG synthesis on the sPG-track^13,14^. However, it is unclear how far this model generalises across the bacterial domain, as many species lack a known activator of cell division.

Here, we investigated the dynamics of the divisome synthesis complex and its coordination with FtsZ in *B. subtilis*. We imaged the divisome transpeptidase PBP2B at a single-molecule level around the entire division septum by orienting cells vertically in bacteria-shaped holes^9,15^. In contrast to the predictions of the coupled-complex and two-track models, we found a single population of processively-moving PBP2B that is dependent on sPG synthesis and not associated with FtsZ treadmilling. Although the motions of PBP2B and FtsZ are asynchronous, we found that the speeds of processive PBP2B molecules are partially dependent on FtsZ treadmilling. Additionally, we provide evidence that the divisome synthesis complex is multimeric. Our results support a new model for division in *B. subtilis* where a multimeric divisome synthesis complex follows a single track dependent on sPG synthesis whose activity and dynamics are asynchronous with FtsZ treadmilling.

## Results

### Processive motion of the divisome synthesis complex depends on sPG synthesis

We created a model of the *B. subtilis* divisome core complex (consisting of the proteins PBP2B, FtsW, FtsL, DivIB, and DivIC) using AlphaFold2 Multimer (Methods; Supplementary Figure 1), showing close agreement with the recent cryo-EM structure of homologous proteins from *Pseudomonas aeruginosa*^6^. Since a recent study showed that these five proteins move together as a complex *in vivo*^7^, we decided to follow the overall motion dynamics of the divisome synthesis complex by tracking the well-characterised transpeptidase PBP2B.

We constructed a strain that expresses a previously-characterized HaloTag (HT) fusion of PBP2B as a functional sole copy at its native locus from an IPTG-inducible promoter^2^ (Methods; Supplementary Table 1). Induction of protein expression with 100 μM IPTG gave near-native cell morphology (Supplementary Figure 2), with HT-PBP2B levels at ∼67% of native PBP2B levels (Supplementary Figure 3). We chose 100 μM IPTG induction of HT-PBP2B for experiments as higher induction levels did not produce significant changes in cell morphology (Supplementary Figure 2). To identify division septa, in addition to unlabelled native FtsZ our strain expresses GFP-FtsZ from an ectopic locus at low levels (0.075% xylose induction; Methods) that do not interfere with cellular growth (Supplementary Figure 4) or FtsZ treadmilling speed (Supplementary Figure 5).

To image single-molecule dynamics of PBP2B around the septum throughout division, we used smVerCINI (single-molecule Vertical Cell Imaging by Nanostructured Immobilisation), a method we developed for single-molecule imaging in rod-shaped cells by confining cells vertically in nanofabricated micro-holes^16^ (Figure 1a). For single-molecule resolution, we sparsely labelled cells with JFX554 HaloTag ligand^17^ (100-250 pM; Methods) and loaded them into agarose micro-holes as described previously^9,15^. We identified division rings using the dilute GFP-FtsZ signal, and then imaged single-molecule dynamics of HaloTag-PBP2B (HT-PBP2B) in rich media at 30°C (Methods; Figure 1b, c; Supplementary Videos 1 and 2).

**Figure 1.**
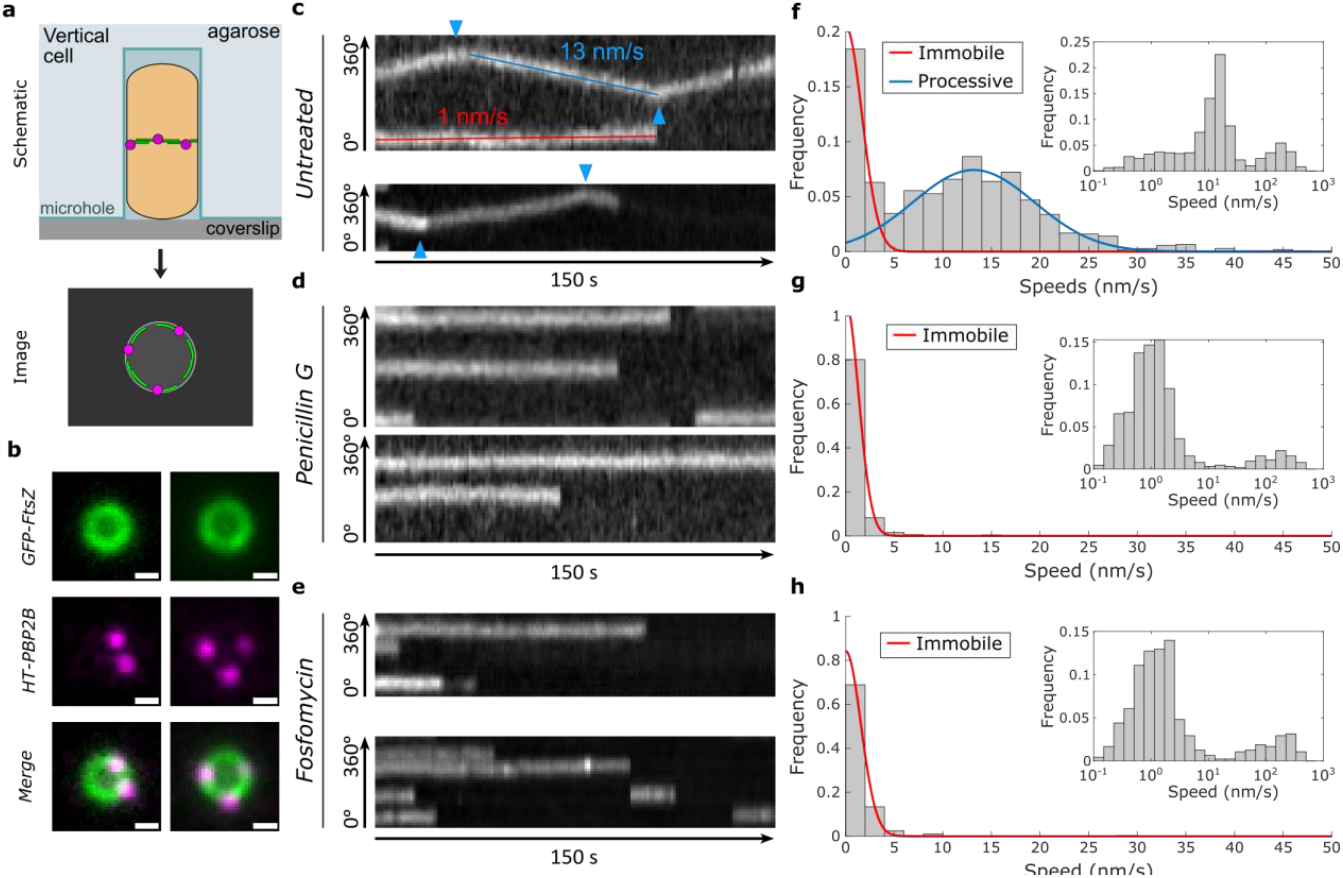
Two-colour smVerCINI imaging shows a single processive population of HT-PBP2B that is associated with sPG synthesis. **(a)** Schematic of two-colour smVerCINI experimental setup. A *B. subtilis* cell (orange) is oriented vertically in a microhole made of agarose (light blue), adjacent to a microscope coverslip (grey). By orienting the cell vertically, the division proteins FtsZ (green) and PBP2B (magenta) can be imaged in a single microscope imaging plane. **(b)** Example images of two vertically trapped cells. GFP-FtsZ (green) is expressed at a low level (0.075% xylose induction) for use as a septal marker, while HT-PBP2B (100 μM IPTG induction) is labelled sub-stoichiometrically with JFX554 HaloTag ligand^17^ (250 pM) so that single molecules can be easily observed. Scale bars: 500 nm. **(c)** Example radial kymographs of HT-PBP2B motion from smVerCINI videos in untreated cells. Diagonal lines result from processive motion (example shown with blue line), while horizontal lines result from lack of motion (example shown with red line). Blue arrows indicate where HT-PBP2B molecules changed direction. **(d, e)** Example radial kymographs of HT-PBP2B motion from smVerCINI videos in cells treated with 20 μg/mL penicillin G or 500 μg/mL fosfomycin. **(f)** Histogram of HT-PBP2B speeds in untreated cells. Red and blue lines show fits to the data (Methods). *Inset*: Histogram of speeds plotted on logarithmic x axis, showing three populations. **(g, h)** Histograms of speeds in cells treated with penicillin G or fosfomycin. Red lines show fits to the data (Methods). *Insets*: Histograms of speeds plotted on logarithmic x axis, showing two populations. All examples in **c-h** are from cells grown in rich media at 30°C.

HT-PBP2B molecules showed several distinct motion behaviours. A distribution of linear track segment speeds reveals three populations: an immobile population (∼0 nm/s), a processive population (∼13 nm/s) (Figure 1f), and a broad fast-moving population (>100 nm/s) that becomes visible when speeds are plotted on a logarithmic scale (Figure 1f inset). Individual HT-PBP2B molecules were also capable of transitioning between states (Supplementary Figure 6). The probabilities of transitioning between states and their associated rates for all conditions in this study are listed in Supplementary Table 2. Based on the measured lifetimes of these states, an HT-PBP2B molecule under our standard conditions exists in the immobile state 38.1 ± 0.4% of the time, the processive state 59.0 ± 0.6%, and the fast-moving state 3.0 ± 0.1% (mean ± SEM; Supplementary Table 3).

We considered that the immobile population could represent molecules bound to the middle of treadmilling FtsZ filaments, as suggested previously for FtsW in *E. coli*^11^. However, the average lifetime of the immobile state of HT-PBP2B molecules (48 ± 3 s (mean ± SEM); Supplementary Figure 7) is substantially longer than the reported lifetime of FtsZ monomers at the division septum in *B. subtilis* (8.1 s)^7^, making this situation unlikely. Due to the long lifetime of this population, we speculate that the immobile population may be bound to the cell wall. It is unclear how such binding would occur, although one attractive possibility is that immobile PBP2B molecules are bound to acceptor peptides in the cell wall awaiting the emergence of nascent glycan strands to crosslink, as proposed recently for the *E. coli* elongasome transpeptidase PBP2^18^.

Processively moving molecules showed a variety of noteworthy behaviours. Individual processive runs were usually terminal (54 ± 4%; mean ± SEM; Supplementary Table 2), indicating that they ended in dissociation, photobleaching, or the end of the acquisition period. However, they often ended with a change in direction (29 ± 3%), or sometimes by becoming immobile (12 ± 2%) or changing to the fast-motion state (5 ± 1%; Supplementary Figure 6). We also observed numerous cases of processive tracks apparently crossing one another (Supplementary Figure 8), suggesting that multiple divisome synthesis complexes exist in different lanes at the septal leading edge, slightly out of plane from one another. The speeds of processive molecules were independent of septal diameter (Supplementary Figure 9), suggesting that their dynamics remain consistent throughout active constriction.

We wondered if the synthesis complex requires sPG synthesis for processive motion. To test this, we imaged the motion of HT-PBP2B immediately after treating cells with an excess (20 μg/mL) of penicillin G, which directly binds the enzyme’s catalytic site and prevents transpeptidation. This abolished the processive population (13.4 nm/s mean; 95% CI [12.7, 14.1]), leaving only immobile and diffusive tracks (Figure 1d, g; Supplementary Videos 3 and 4). We repeated the experiment with fosfomycin, a separate class of antibiotic that inhibits the synthesis of lipid II, the substrate for cell wall synthesis^19^. As with penicillin G, immediately after treatment with an excess (500 μg/mL) of fosfomycin, the processive population of HT-PBP2B vanished (Figure 1e, h; Supplementary Videos 5 and 6). We conclude that processive motion of the synthesis complex requires sPG synthesis.

We next wondered what the broad high speed population was. This population resulted from short back-and-forth tracks that appeared to show diffusive rather than processive motion (Supplementary Figure 10a). These tracks were often located at larger radii than immobile or processive tracks (Supplementary Figure 10b) and were nearly the only type of motion pattern outside the septal ring area (Supplementary Figure 11), suggesting that they are not typically present at the septal leading edge and therefore unlikely to be involved in sPG synthesis. We were initially surprised to find an apparently diffusive population during our experiments, as we expected diffusive motion to be fast enough to be blurred out during our long (1 s) acquisition interval. Since linear speeds did not represent this population well, we instead measured the mean-squared displacements (MSDs) of these tracks and fitted an anomalous diffusion model to them (Methods; Supplementary Figure 10c-f). The effective diffusion coefficient (*D*_*eff*_ = 6.5×10^-3^ ± 0.4×10^-3^ μm^2^/s (mean ± SEM)) and diffusion exponent (*α* = 0.80 ± 0.03 (mean ± SEM)) we obtained indicate that these tracks represent very slow sub-diffusive behaviour. The low diffusion coefficient is similar to that previously measured for the *E. coli* elongasome transpeptidase PBP2^18^, and suggests that diffusive HT-PBP2B molecules may principally exist as part of large multi-protein complexes (i.e. the divisiome core complex). Alternatively, diffusive HT-PBP2B molecules may experience substantial molecular friction through transient interactions with the cell wall.

### The divisome synthesis complex and FtsZ move asynchronously

The speed distributions for HT-PBP2B we measured (Figure 1) are not consistent with either the coupled-complex^2^ or two-track^11^ models. Both models predict that there exists at least one processive population of synthesis complexes that move at the same speed as treadmilling FtsZ filaments^2,11^, but the speed of the processive HT-PBP2B population (13.4 nm/s mean; 95% CI [12.7, 14.1]) (Figure 1f) does not match what we previously measured for FtsZ treadmilling (44.1 nm/s median; 95% CI [26.0, 53.9])^9^ under identical growth conditions (rich media, 30°C). This result does not depend on protein expression levels, as inducing HT-PBP2B with a 10-fold higher concentration of IPTG resulted in similar speeds for processive HT-PBP2B molecules (Supplementary Figure 12). Furthermore, in contrast to predictions from the two-track model^11^, we do not observe the emergence of an FtsZ-associated speed population of HT-PBP2B after treating cells with penicillin G or fosfomycin (Figure 1g, h).

Next, we investigated whether the asynchronous motions of synthesis complexes and FtsZ treadmilling are a general feature in *B. subtilis* by repeating our measurements under different growth conditions (Figure 2a). Our results show that HT-PBP2B speeds depended on both media composition and temperature: processive molecules moved faster in rich media or 37°C than in minimal media or 30°C (blue lines in Figure 2a). Under most growth conditions tested, the speeds of processive HT-PBP2B molecules do not match the speeds of FtsZ treadmilling measured under identical conditions^9^ (green dotted lines in Figure 2a). However, under the fastest growth condition (rich media, 37°C) HT-PBP2B processive speeds overlap substantially with FtsZ treadmilling speeds. Notably, this was the growth condition under which single-molecule tracking of HT-PBP2B was performed previously^2^. The similarity of speeds between FtsZ treadmilling and HT-PBP2B processive motion in those conditions led us to initially propose that the two systems moved together as a coupled complex. Our results instead indicate that this overlap in speeds is a coincidence arising only under the fastest growth condition (Figure 2a).

**Figure 2.**
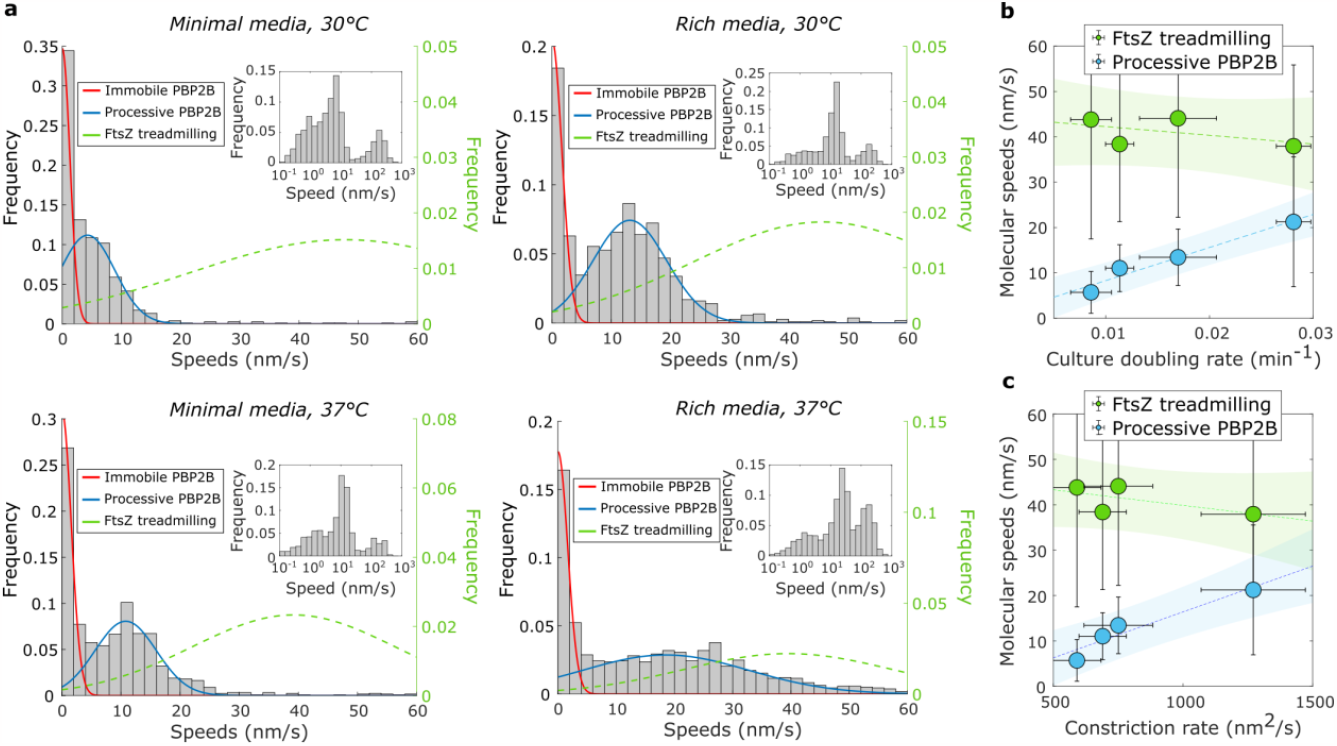
PBP2B and FtsZ move asynchronously. **(a)** Histograms of HT-PBP2B speeds across growth conditions. Red and blue lines show fits to the data (Methods). Green dotted lines show Gaussian distributions based on the mean and variance of FtsZ treadmilling speeds measured under the same growth conditions (data from Whitley et al. 2021^9^). *Insets*: Histograms of speeds plotted on logarithmic x axes, showing three populations. **(b)** Correlation between culture doubling rates with speeds of FtsZ treadmilling and processive HT-PBP2B. Dotted lines show linear fits to the data. Shaded regions show 95% confidence bands. Error bars are SD. **(c)** Correlation between FtsZ treadmilling and processive HT-PBP2B speeds with septal constriction rate. Septal constriction rates under these conditions are from Whitley et al. 2021^9^. Dotted line shows a linear fit to the data. Shaded region shows 95% confidence bands. FtsZ treadmilling speeds: median ± SD. HT-PBP2B speeds: mean ± SD.

The variance of the processive HT-PBP2B population under the fastest growth condition is substantially higher than that of other conditions (Figure 2a). The reason for this increase is unclear, although it was reproducible: each of the four biological replicates we performed under this condition had a similarly large variance. This increase is larger than the dependence predicted by a Poisson process, which is commonly observed in single molecule dynamics. Furthermore, it does not seem to result from perturbed FtsZ treadmilling in the particular strain used here, as the speed distribution for FtsZ treadmilling under these conditions was comparable to those of previous measurements (Supplementary Figure 5). It is possible that the increased variance in HT-PBP2B speeds under this condition reflects increased variation in the local production or local availability of the lipid II substrate used by the divisome synthesis complex. However, this is currently difficult to investigate experimentally.

We found that processive HT-PBP2B speeds are correlated with cell growth rate (doubling rate in liquid culture), while FtsZ treadmilling is independent of growth rate (Figure 2b). We speculate that faster growth conditions results in more rapid production of cell wall substrate (lipid II), leading to a higher rate of sPG incorporation reactions and hence higher synthesis complex speeds. We further found that processive HT-PBP2B speeds are correlated with the rate of septal constriction, which we measured previously under identical conditions^9^ (Figure 2c). It is possible that higher synthesis complex speeds lead to more sPG added to the ingrowing septum per unit time, thereby resulting in a higher rate of septal constriction.

It is surprising that FtsZ treadmilling speed is independent of temperature, as it depends on an enzymatic reaction. Using the Eyring equation with activation enthalpies measured previously for *E. coli* FtsZ *in vitro*^20^, we predict that an increase of 30°C to 37°C should result in a ∼2-fold increase in treadmilling speed (Supplementary Note 2), which we have not observed. To test how far this temperature independence extrapolates, we measured FtsZ treadmilling speed at 21°C using a previously characterized strain (bWM4; Supplementary Table 1) expressing mNeonGreen-FtsZ from an IPTG-inducible promoter (Supplementary Figure 13), grown in rich media. Treadmilling speed was 32% slower than it was at 30°C, although this is a substantially smaller decrease than the 68% predicted from the Eyring equation. This suggests that bacterial cells may actively regulate the polymerization dynamics of FtsZ filaments, as the lack of temperature dependence of treadmilling speed is not explained by chemical physics alone.

### Processive motion of the divisome synthesis complex partially depends on FtsZ treadmilling

Our finding that FtsZ and HT-PBP2B move asynchronously suggests that synthesis complexes are independent of FtsZ treadmilling in *B. subtilis*. To test this directly, we measured the dynamics of HT-PBP2B in the absence of FtsZ treadmilling. We treated cells with 10 μM PC190723, an antibiotic that specifically binds to FtsZ^21^ and arrests treadmilling^2^ across all stages of division within seconds of treatment^9^. HT-PBP2B still showed processive motion (Figure 3a, c; Supplementary Videos 7 and 8), but the speeds were slower than in untreated cells (8.1 nm/s mean; 95% CI [7.0, 9.4] treated vs. 13.4 nm/s mean; 95% CI [12.7, 14.0] untreated). This is a stark difference from the penicillin G-or fosfomycin-treated cells, where the processive population was abolished (Figure 1g, h). We confirmed that sPG synthesis continues in the absence of FtsZ treadmilling by treating cells with PC190723 for approximately one round of cell division while simultaneously labelling them with fluorescent D-amino acids, then imaging with Structured Illumination Microscopy (Methods; Supplementary Figure 14).

**Figure 3.**
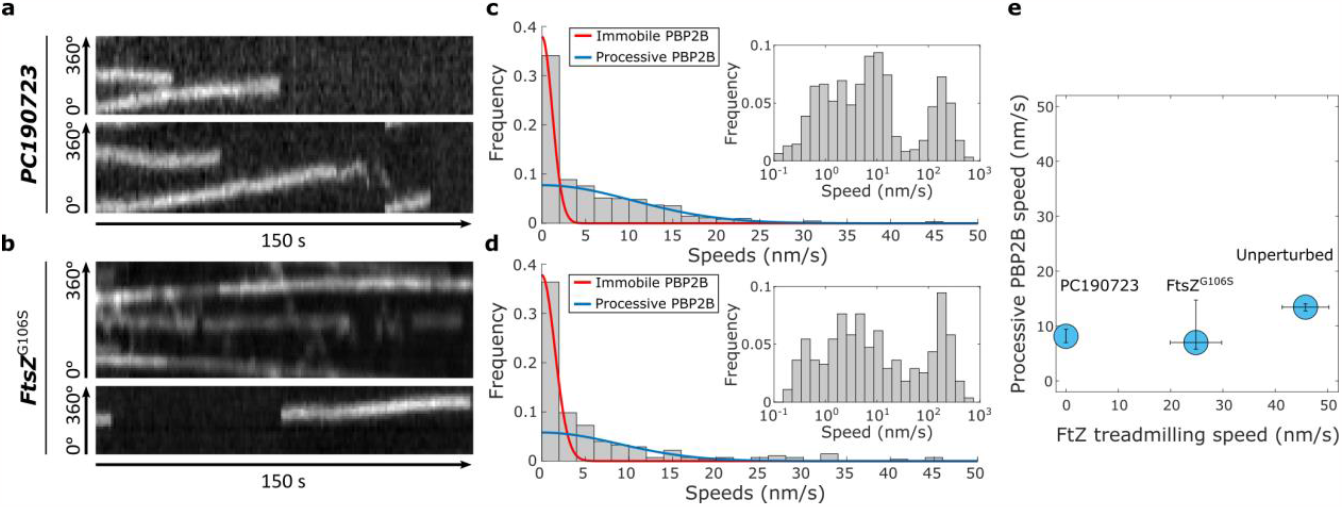
PBP2B motion partially depends on FtsZ treadmilling. **(a)** Example radial kymographs of HT-PBP2B motion from smVerCINI videos in cells treated with 10 μM PC190723. **(b)** Example radial kymographs in cells expressing the treadmilling-impaired mutant FtsZ^G106S^. **(c)** Histogram of speeds in cells treated with PC190723. **(d)** Histogram of speeds in FtsZ^G106S^ cells. Red and blue lines to both histograms show fits to the data (Methods). *Insets*: Histograms of speeds plotted on logarithmic x axes, showing three populations. **(e)** Relation between FtsZ treadmilling speeds and processive HT-PBP2B speeds. The value for FtsZ^G106S^ treadmilling speed was measured in this study under our experimental conditions (Supplementary Figure 15). Values shown are mean ± 95% CI.

We wondered if this drop in speeds was specific to treatment with PC190723, which is a strong perturbation due to its sudden and total arrest of FtsZ treadmilling. We therefore tried a similar genetic perturbation by using an FtsZ mutant (FtsZ^G106S^) that is competent for division but has reduced treadmilling speed^2^ (Supplementary Figure 15) and produces a long-cell phenotype (Methods; Supplementary Table 1; Supplementary Figure 16). The speed distribution for HT-PBP2B in this mutant strain was similar to that observed with PC190723-treated wild-type cells (Figure 3b, d; Supplementary Videos 9 and 10). This suggests that any substantial disruption to FtsZ treadmilling has similar effects on synthesis complex motion (Figure 3e), in contrast to the linear relation between their speeds proposed previously^2^.

We recently reported evidence from computational studies that perturbations to FtsZ treadmilling disrupt the nematic order of the FtsZ filament network^22^. We therefore considered that the reduction in HT-PBP2B speeds could result from motion along a disordered, jagged path due to transient interactions between the divisome synthesis complex and randomly-oriented FtsAZ filaments. To test this, we imaged the processive motion of HT-PBP2B molecules in horizontally-oriented cells and measured the displacements from the septal axis (Methods; Supplementary Videos 14-16; Supplementary Figure 17). With either PC190723 treatment or expression of FtsZ^G106S^, the median off-axis displacements were within 2 nm of that measured for the unperturbed case, which is unlikely to be biologically meaningful (Supplementary Figure 17e). This suggests that the reduction in HT-PBP2B speeds observed upon perturbations to FtsZ treadmilling does not result from HT-PBP2B off-axis motion.

### The divisome synthesis complex is multimeric

During this study, we observed many cases where the fluorescence intensities of HT-PBP2B spots showed discrete drops to half their value (Figure 4a, b; Supplementary Videos 2 and 11), indicating the presence of two copies of the fluorescently-labelled protein. Under our standard conditions (rich media, 30°C, 100 μM IPTG, 250 pM JFX554 HaloTag ligand), we observed these intensity drops in 11% (N=56) of full tracks. We also observed multiple occasions where such intensity drops occurred during motion and even after direction changes (Figure 4a, b; Supplementary Figure 18), strongly suggesting that multiple monomers of HT-PBP2B are moving together as part of a larger complex. Due to the sub-stoichiometric nature of the labelling method, we cannot precisely quantify the number of HT-PBP2B molecules in a given complex, although we have observed rare cases with even three or four such drops in intensity (Supplementary Figure 19).

**Figure 4.**
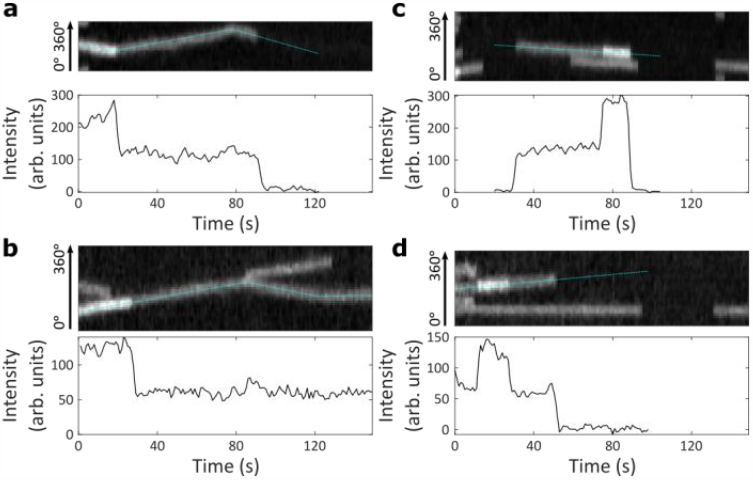
PBP2B is part of dynamic multimeric complexes. **(a, b)** *Top:* Example kymographs of HT-PBP2B motion from smVerCINI videos showing discrete drops in fluorescence intensity mid-track. *Bottom:* Intensity traces for the tracks designated by dotted cyan lines overlaid on kymographs. Both traces are from cells grown in rich media at 30°C. **(c, d)** *Top:* Example kymographs showing discrete jumps in fluorescence intensity mid-track. *Bottom:* Intensity traces for the tracks designated by dotted cyan lines overlaid on kymographs. Both traces are from cells grown in minimal media at 30°C.

Surprisingly, we also observe cases where the fluorescence intensity signal shows discrete jumps to twice their value (4% (N=22) of full tracks under our standard conditions; Figure 4c, d; Supplementary Figure 20; Supplementary Videos 12 and 13). We observe these discrete jumps in both immobile and processive tracks. This suggests that the oligomeric state of synthesis complexes is dynamic, where new PBP2B molecules can bind to both active and inactive complexes. As only 3 out of 22 observed intensity jumps (14%) and 9 out of 56 (16%) of observed intensity drops roughly corresponded to a change in HT-PBP2B speed, it appears that these events do not necessarily affect divisome synthesis complex activity, although it remains possible that there is a higher probability of speed change during an intensity drop/jump event than without.

We wondered whether this behaviour was unique to PBP2B, or if it was a more general feature of divisome synthesis complex proteins. We repeated these measurements with a strain expressing a HaloTag fusion of the transglycosylase FtsW (HT-FtsW; Supplementary Table 1) that together with PBP2B forms the core of the synthesis complex. HT-FtsW displayed similar fluorescence drops and jumps as HT-PBP2B (Supplementary Figure 21), suggesting that the divisome synthesis complex is multimeric. The effect of stoichiometry on divisome activity and dynamics will be followed up in future work.

## Discussion

Our results show that a multimeric divisome synthesis complex in *B. subtilis* follows a single track dependent on sPG synthesis and asynchronous with FtsZ treadmilling. This sharply contrasts with two prominent models for bacterial cell division^2,11^, which predict the existence of a processive population of synthesis complexes associated with FtsZ treadmilling. Our results instead support a model of septal PG synthesis where the Z-ring recruits the synthesis complex to the septal leading edge but does not directly regulate its motion and synthesis activity (Figure 5).

**Figure 5.**
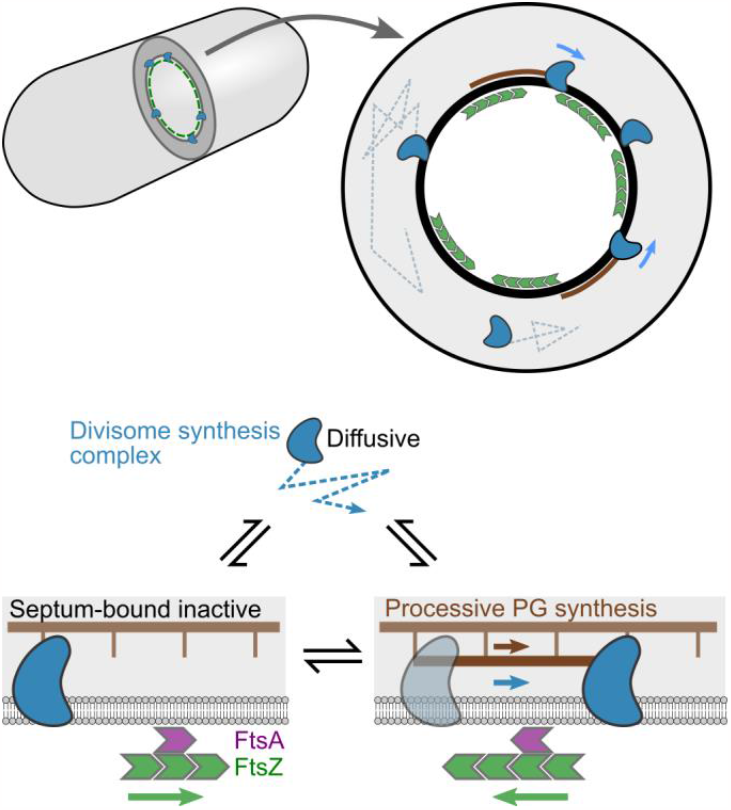
A one-track model for spatiotemporal regulation of *B. subtilis* septal peptidoglycan synthesis activity. Inactive divisome synthesis complexes (blue) diffuse around the cell membrane and are recruited to the septal leading edge, marked by a dense FtsZ-ring (green), via transient interactions with the FtsZ anchors FtsA (purple) and SepF (not shown for clarity). Once recruited to the septum, synthesis complexes move asynchronously from FtsZ, switching between an inactive static state, potentially bound to the cell wall, and an active state performing processive septal PG synthesis

Our finding that the motions of synthesis complexes and FtsZ are asynchronous is consistent with previous measurements in *S. pneumoniae*^5^ and work performed in parallel to this study in *S. aureus*^23^. Our discovery that synthesis complexes retain processive motion in the absence of FtsZ treadmilling also agrees with previous findings in both *S. aureus* and *B. subtilis* that sPG synthesis continues in the absence of FtsZ treadmilling if constriction has already initiated^4,9^. This suggests that these species—all members of the phylum Bacillota (Firmicutes)—share a similar mechanism for division. It is possible that the two-track model proposed for *E. coli* and *C. crescentus*—both members of the phylum Pseudomonadota (Proteobacteria)—represents a special case arising from requirements unique to this bacterial clade, such as a thinner PG layer or sparser network of FtsZ filaments.

Previously, our collaborators observed that arrest of FtsZ treadmilling by the antibiotic PC190723 abolished processive HT-PBP2B motion^2^, but here we find that the effect of arresting FtsZ treadmilling is to slow HT-PBP2B processive motion rather than stop it. One difference between these studies is that cells were imaged here with VerCINI while the previous study imaged horizontally-oriented cells with Total Internal Reflection Fluorescence (TIRF) illumination. However, as part of this study we also imaged horizontally-oriented cells with TIRF and observed processive motion at division septa post-PC190723 treatment (Supplementary Video 15). It is possible that processive motion was not observed in the previous study due to small sample size.

We observed that perturbations to FtsZ treadmilling lead to slower speeds for processive synthesis complexes. The reduction in speeds upon total arrest of FtsZ treadmilling (∼40%) correlates with the reduction in septal constriction rate (by 14-33%, depending on growth conditions) that we previously observed under similar conditions^9^. This is corroborated by our finding that synthesis complex speeds are correlated with septal constriction rate (Figure 2c). However, work performed in parallel to this study in *S. aureus* found that synthesis complex speeds and septal constriction rate are not affected by FtsZ treadmilling^23^. In contrast to both of these studies, previous work in *S. pneumoniae* found a minor (∼25%) reduction in synthesis complex speeds when FtsZ treadmilling was severely perturbed, and no effect from a smaller perturbation^5^. This suggests that although these three species may share a similar mechanism for division, there may remain differences, which could arise from changes in the interaction strength of FtsZ subunits with the rest of the divisome.

The underlying mechanism for the reduction in synthase complex speeds upon FtsZ treadmilling perturbation remains unclear. It is possible that the disruption to FtsZ treadmilling leads to an alteration of transient interactions between FtsAZ filaments and the divisome synthesis complex, leading to increased molecular friction. However, we consider this unlikely, as both a reduction and total arrest of FtsZ treadmilling produced similar effects on HT-PBP2B speeds (Figure 3e). The reduction in HT-PBP2B speeds may instead be an indirect effect of disrupting FtsZ treadmilling. We have shown that the speeds of the divisome synthesis complex depends on cell metabolism (Figure 2b). It is plausible that severe perturbations to an essential and abundant protein such as FtsZ could affect metabolism (e.g. through a stress response) and hence indirectly reduce synthase speeds.

Our results, along with those in multiple other organisms^5,23^, strongly support a model where the processive motion of sPG synthases in Bacillota is driven exclusively by sPG synthesis. Our observation that treatment with antibiotics targeting either synthase activity or lipid II precursor synthesis prevents processive motion (Figure 1c-h) suggests that the sPG insertion reaction itself may provide the required energy. A similar mechanism was previously proposed for elongasome synthases^24–26^. Further work is required to understand the molecular mechanism by which SEDS-bPBP PG synthesis activity leads to processive motion.

We found evidence that the divisome synthesis complex—the motile multi-protein complexes of divisome proteins that move around the septum synthesising PG—is multimeric, in support of the stoichiometric divisome hypothesis^27^. In *Pseudomonas aeruginosa*, the structure of the so-called divisome core complex of proteins, which is likely the minimum holoenzyme unit required to synthesise and attach a single strand of septal PG, was reported^6^. We found that AlphaFold predicts a *Bacillus subtilis* divisome core complex very similar to the recently published structure of the *Pseudomonas aeruginosa* divisome core complex (Supplementary Figure 1), consisting of PBP2B (*P. aeruginosa* FtsI/ PBP3 homologue) in a complex with FtsW, DivIC (FtsB homologue), FtsL and DivIB (FtsQ homologue). As the stoichiometry of PBP2B and FtsW with each of the other components in the divisome core complex is 1:1, our data suggest that individual divisome synthesis complexes often contain multiple divisome core complexes. The multimeric nature of the divisome synthesis complex may also explain the abrupt changes in direction we observe in single-molecule tracks (Figure 1c). Such bidirectional motion could result from multiple synthesis proteins in a complex pulling in opposite directions, as is well-known for motor proteins moving along the eukaryotic cytoskeleton^28^ and more recently the elongasome of *B. subtilis*^16^. Inclusion in a large multimeric complex may also explain the very low effective diffusion coefficient we measured for HT-PBP2B in the septal ring (Supplementary Figure 10e). These possibilities will be investigated in future studies.

## Acknowledgements

We would like to acknowledge Ethan Garner (Harvard) for sharing strains, Ling Juan Wu (Newcastle) for sharing strains and antibodies, Luke Lavis (Janelia Farm) for providing us with JFX554 HaloTag ligand and Stuart Middlemiss (Newcastle) for advice with cloning. We would also like to thank Mariana Pinho (ITQB NOVA) and Jie Xiao (Johns Hopkins) for helpful discussions. S.H., K.D.W., D.M.R., J.G., and E.K. acknowledge funding support by a Newcastle University Research Fellowship, a Wellcome Society Sir Henry Dale Fellowship grant number [206670/Z/17/Z], and a BBSRC 19ALERT mid-range equipment initiative grant [BB/T017570/1]. Research in P.J.S.’s lab was funded by Wellcome (208361/Z/17/Z), the MRC (MR/S009213/1) and BBSRC (BB/P01948X/1, BB/R002517/1 and BB/S003339/1). We acknowledge use of Zeiss Lattice SIM2 within the School of Life Sciences Microscopy Facility, University of Warwick and a BBSRC ALERT grant BB/W020300/1 to S.H. which supported purchase of that microscope.

## Author contributions

K.D.W. and S.H. designed the research; K.D.W. and D.M.R. performed the experiments; K.D.W., J.G., and E.K. constructed and characterised bacterial strains; K.D.W., D.M.R., and J.G. analysed the data. P.J.S. performed protein structure predictions; K.D.W. and S.H. wrote the manuscript with input from all authors.

## Methods

### Divisome complex modelling

The divisome complex was modelled by AlphaFold2, using ColabFold (v1.3.0) and AlphaFold-Multimer (v2). The sequences were downloaded from the UniProtKB database and included five divisome protein sequences (Q07868 (PBP2B); Q07867 (FtsL); O07639 (FtsW); P16655 (DivIB); P37471 (DivIC)).

### Bacterial strains and growth conditions

Strains used in this study are listed in Supplementary Table 1. Strains were streaked from -80°C glycerol stocks onto nutrient agar (NA) plates containing the relevant antibiotics and 1 mM IPTG and grown overnight at 37°C. Single colonies were transferred to liquid starter cultures in either PHMM medium^2^ or Time-Lapse Medium^29^ (TLM) with required inducers, shaking at 200 rpm overnight at either 22°C or 30°C. The next day, PHMM starter cultures were diluted to OD_600_ = 0.05 in fresh PHMM, while TLM starter cultures were diluted OD_600_ = 0.1 in Chemically-Defined Medium (CDM)^29^. These liquid cultures were grown at 30 or 37°C with required inducers until they reached 0.3 < OD_600_ < 0.6. When necessary, antibiotics were used at the following concentrations: spectinomycin 60 μg/mL, erythromycin 1 μg/mL, lincomycin 25 μg/mL, 6 μg/mL tetracycline.

### Strain construction

Strains SH147 and SH203 harboured a deletion of the *hag* gene, as this has been shown to reduce the chaining phenotype in *B. subtilis* cells and thereby increase loading into micro-holes^9,15^.

SH142 (PY79 Δ*hag, amyE*::*spc-*P_xyl_*-gfp-ftsZ*) was constructed by transforming SH211 with genomic DNA extracted from strain 2020 using standard protocols^30^.

SH147 (PY79 Δ*hag, pbpB*::*erm-*P_hyperspank_-*HaloTag-15aa-pbpB, amyE*::*spc-*P_xyl_-*gfp-ftsZ*) was constructed by transforming SH142 with a PCR product obtained from bGS31 genomic DNA. The primer pair ftsL Fw/ spoVD Rev (Supplementary Table 4) were used to amplify the region around the *pbpB* gene using bGS31 genomic DNA as a template. The primer pair ftsL Fwd/ pbp2B Rev (Supplementary Table 4) was used to confirm insertion of the *HaloTag* gene in the transformant. Insertion was also confirmed by Sanger sequencing.

SH203 (PY79 Δ*hag, pbpB*::*erm*-P_hyperspank_-*HaloTag-15aa-pbpB, ftsZ*Ω*ftsZ(G106S) (tet), amyE*::*spc-*P_xyl_-*gfp-ftsZ*) was constructed by transforming SH147 with genomic DNA extracted from strain Z-G106S (gifted by Ethan Garner). The point mutation G106S was confirmed by Sanger sequencing.

All published strains are available on request to the authors.

### Bacterial strain characterization

Strains were characterised by growth in liquid culture (Supplementary Figures 4 and 16) and cell morphology analysis (Supplementary Figures 2 and 16).

### Growth curves

*B. subtilis* PY79 and variant strains were grown in liquid starter cultures overnight in LB at 30°C with required inducers (100 μM IPTG for SH147 and SH203). To measure culture growth across [IPTG] (Supplementary Figure 4), SH147 overnight cultures were washed twice in LB to remove inducer. Cultures were then diluted to OD_600_ = 0.05 in LB with variable inducer concentrations, and 200 μL of each dilution was added to a 96-well microtiter plate. Growth was monitored for 15 h using a FLUOStar OPTIMA plate reader (BMG Labtech). Growth curves for each condition were performed in triplicate.

### Western blotting

Overnight cultures of specified strains were grown overnight in PHMM at 22°C. The following morning, cultures were diluted to OD_600_∼0.05 and grown at 37°C until OD_600_∼0.4. Cells were harvested by centrifugation and lysed by incubation for 20 min in BugBuster protein extraction reagent supplemented with Benzonase nuclease (Millipore) and an EDTA free protease inhibitor cocktail (Roche). Protein extract was heated for 10 min at 65°C in NuPAGE LDS Sample Buffer, then 5 μg total protein in this buffer was separated by SDS-PAGE on a NuPAGE 3-8% Tris-Acetate Midi Gel (Invitrogen). Protein was transferred to a 0.45 μm PVDF Membrane (Cytiva), and PBP2B and Spo0J were detected using PBP2B polyclonal and Spo0J polyclonal antibodies, respectively, followed by a HRP-conjugated anti-rabbit IgG antibody (Sigma). Samples were developed using Clarity Western ECL Substrate (Bio-Rad) and imaged using an ImageQuant LAS 4000 mini Biomolecular Imager (GE Healthcare).

### Cell morphology analysis

*B. subtilis* Δ*hag* (strain SH211) and variant strains were grown in liquid starter cultures overnight in PHMM at 22°C or 30°C with required inducers (100 μM IPTG for SH147 and SH203). To measure cell morphology across [IPTG] (Supplementary Figure 2c), SH147 overnight cultures were washed twice in PHMM to remove inducer. Cultures were then diluted to OD_600_ = 0.05 in PHMM with variable [IPTG] and 0.075% xylose. To measure cell morphology across [xylose] (Supplementary Figure 2d), overnight SH147 cultures were diluted to OD_600_ = 0.05 in PHMM with 100 μM IPTG and variable [xylose]. Once the cultures had reached 0.3 < OD_600_ < 0.6, Nile Red was added to cells and incubated at growth temperatures for 5 min. 0.5 μL of cell culture was then spotted on 1.2-2% agarose pads of PHMM with the required inducers, prepared as described previously^29^. Cells were imaged using a 561 nm laser or 550 nm LED, and cell lengths were manually determined using ImageJ.

### smVerCINI

smVerCINI was set up as described previously for VerCINI^9,15^. Briefly, agarose microholes were formed by pouring molten 6% agarose onto a nanofabricated silicon array consisting of micropillars with widths 1.0-1.3 μm and heights 6.8 μm. Patterned agarose was transferred into a Geneframe (Thermo Scientific) mounted on a glass slide, and excess agarose was cut away to ensure sufficient oxygen. Labelled cells at 0.3 < OD_600_ < 0.6 were concentrated 100× by centrifugation and added onto the agarose pad. Cells were then loaded into the microholes by centrifuging the mounted agarose pad with concentrated cell culture in an Eppendorf 5810 centrifuge with MTP/Flex buckets.

Unloaded cells were rinsed off with excess media. In experiments where cells were treated with antibiotic (Figures 1d, 1e, and 3a), 5 μL of media laced with antibiotic was added to the top of loaded cells and allowed to absorb for ∼1 min prior to sealing with a coverslip and imaging.

Imaging was done by first recording a single frame of GFP-FtsZ using the 488 nm laser to identify division rings. Immediately following this, a time-lapse of HT-PBP2B dynamics was recorded using the 561 nm laser. Following fluorescence imaging, a short bright-field video was recorded to identify any cells that were improperly trapped in micro-holes. Microscopy acquisition parameters are listed in Supplementary Table 5.

### HaloTag labelling with JFX554 dye

Single-molecule labelling of HT-PBP2B was done by incubating strain SH147 or SH203 with either 100 pM (minimal media) or 250 pM (rich media) JFX554 HaloTag ligand for 15 min unless otherwise noted. Cells were washed once with fresh media before imaging. JFX554 HaloTag ligand was a gift from Luke Lavis (Janelia Farm)^17^.

### Microscopy

Power densities, exposure times, and other key parameters are listed for each microscopy experiment in Supplementary Table 5.

#### Nikon Eclipse Ti2

Cells were illuminated with 488 nm and 561 nm laser illumination. A 100× TIRF objective (Nikon CFI Apochromat TIRF 100XC Oil) was used for imaging and a Kinetix sCMOS camera (Teledyne Photometrics) was used with effective pixel size of 65 nm/pixel. Cells were illuminated using HiLO or TIRF to minimise background using an objective TIRF module.

#### Bespoke microscope

Cells were illuminated with a 488 nm laser (Obis) and a 561 nm laser (Obis). A 100× TIRF objective (Nikon CFI Apochromat TIRF 100XC Oil) was used for all experiments. A 200 mm tube lens (Thorlabs TTL200) and Prime BSI sCMOS camera (Teledyne Photometrics) were used for imaging, giving an effective pixel size of 65 nm/pixel. Imaging was done with a custom-built ring-TIRF module operated in ring-HiLO using a pair of galvanometer mirrors (Thorlabs) spinning at 200 Hz to provide uniform, high SNR illumination^31^.

#### Nikon Eclipse Ti

Cells were illuminated with a 550 nm LED (CoolLED). A 100× TIRF objective (Nikon Plan Apo 100×/1.40 NA Oil Ph3) was used with a Prime BSI camera (Teledyne Photometrics).

#### Zeiss Elyra 7 Lattice SIM2

Cells were illuminated with 488 nm and 561 nm lasers. A 63× objective (Plan Apo 63x/1.40 Oil) was used for SIM experiments. A 1.6x Optovar and two PCO.edge 4.2 sCMOS cameras (PCO Imaging) were used for imaging, giving an effective pixel size of 62 nm/pixel.

### TIRF microscopy of horizontal cells

Coverslips were first cleaned by treating with air plasma for 5 min. Slides were prepared as described previously^29^. Flat 2% agarose pads of PHMM containing inducers were prepared inside Geneframes (Thermo Scientific) and cut down to strips of ∼5 mm width to ensure sufficient oxygen supply to cells. Cell cultures were grown to OD_600_ between 0.4 and 0.7, when 0.5 μL of cell culture was spotted on the pad. Cells were allowed to adsorb to the pad for ∼1 min. In the case where the HT-PBP2B GFP-FtsZ Δhag strain (SH147) was treated with PC190723, 1 μL of a solution of PHMM + 100 μM IPTG + 0.075% xylose + 10 μM PC190723 was then spotted on top of the cells and allowed to absorb into the pad for ∼1 min. A plasma-treated coverslip was then placed on top. Cells were allowed to equilibrate within the microscope body for ∼2 min before being imaged. Cells were then imaged using TIRF microscopy to observe either FtsZ treadmilling dynamics or HT-PBP2B dynamics. If the concentration of labelled FtsZ was too high to measure treadmilling speeds, the illumination mode was changed to HiLO for 1-10 s to photobleach the label down to an acceptable level before data was acquired. Experimental parameters are defined in Supplementary Table 5.

### Structured Illumination Microscopy (SIM) of cells labelled with fluorescent D-amino acids

*B. subtilis* PY79 and variant strains were grown in liquid starter cultures overnight in PHMM at 30°C with required inducers (100 μM IPTG for SH147 and SH203). The overnight cultures were then diluted to OD_600_ = 0.1 into fresh PHMM and grown at 30°C until 0.4 < OD_600_ < 0.6. At this point, cultures were re-diluted to OD_600_ = 0.1 in pre-warmed PHMM, and 200 μL of diluted culture was transferred to 2 mL tubes with holes in the lid for aeration. The green fluorescent D-amino acid BADA^32^ (Tocris Bioscience) was added to a final concentration of 0.5 mM and tubes were incubated at 30°C with shaking for 90 min. Samples were washed with 200 μL pre-warmed PHMM. After the second wash, the red fluorescent D-amino acid TADA^33^ (Tocris) was added to a final concentration of 0.5 mM. Where cells were treated with PC190723, this compound was also added to a final concentration of 14 μM. Samples were then re-incubated at 30°C with shaking for 10 min. Cells were then washed once with pre-warmed PHMM prior to the addition of 100% ice-cold ethanol. Samples were fixed on ice for 1 hr. Fixed cells were collected by centrifugation and washed twice with cold phosphate buffered saline (PBS).

0.5 μL cells were spotted onto 2% agarose pads in Geneframes (Thermo Scientific) and allowed to adsorb for several minutes prior to the addition of a coverslip. Cells were imaged by 2D SIM on an Elyra 7 Lattice SIM2 microscope (Zeiss). All images were acquired with an exposure time of 118 ms and a laser power of 2% in each channel. Alignment of the two imaging channels was conducted using Tetraspeck fluorescent beads as fiducial markers (Invitrogen). SIM image processing was performed in Zen Black, in 2D SIM mode, using the Standard configuration. Image registration to correct for residual misalignment between the two imaging channels was performed using Zen Black Channel Alignment tool, fitting an Affine transform between the two imaging channels based on image similarity and then applying it to the red image channel. After SIM reconstruction, images had an effective pixel size of 31 nm.

### Image processing and analysis

Videos were denoised using the GPU-accelerated ImageJ plugin PureDenoise-GPU^16^ or the CPU-based version PureDenoise-CPU^34^.

Cells with mature or constricting division rings in focus were chosen using the first GFP-FtsZ image acquired in the imaging sequence described above. The short bright-field videos of each field of view acquired after fluorescence imaging were then used to filter out any of these cells that were improperly trapped in the holes.

Previously developed software^9,15^ was used to subtract the cytoplasmic background signal and produce radial kymographs. Due to chromatic aberration, the HT-PBP2B signal itself was used for fitting and extracting radial kymographs rather than the single GFP-FtsZ frame. Due to the difficulty of fitting a circle to these sparsely-labelled rings, the maximum intensity projections of the HT-PBP2B videos were used for fitting, and each fit was manually inspected to confirm it was adequate.

Measurements of filament speed and processivity were performed manually by kymograph analysis, annotating filaments as lines in ImageJ and then measuring the angle via ImageJ script. For visual display purposes, kymographs represented here have the circle origin (0°/360°) rotated around the division ring so that single-molecule tracks are appropriately shown as continuous.

### Speed distribution analysis

Speed histograms were fitted to a sum of Gaussian distributions. For most histograms, a sum of three Gaussian distributions was used, but in the cases of penicillin G and fosfomycin treatment (Figure 1g, h) these fits yielded two overlapping distributions at 0 nm/s. In these two cases, then, a sum of two Gaussian distributions was used rather than three. As one population was expected to represent fully immobile molecules with speed of 0 nm/s, this one parameter was fixed in all cases. Processive HT-PBP2B speeds were determined from these fits by calculating the first moment of the Gaussian distribution fitting the processive population. 95% confidence intervals were obtained by bootstrapping.

### Single-particle tracking and MSD analysis

For analysis of the diffusive HT-PBP2B population, single molecules were detected and tracked using TrackMate^35^ with linking distance of 0.5 μm, 5 frame gaps, 0.5 μm gap-closing distance. Only tracks >10 frames long were used for further analysis. Bespoke Matlab code was used to plot tracks in polar coordinates and fit mean-squared displacements (MSDs) to the anomalous diffusion model *MSD* = 4*D*_*eff*_*t*^*α*^, where *D*_*eff*_ is the effective diffusion coefficient and *α* is the anomalous diffusion exponent.

### Statistics and reproducibility

All sample sizes and number of experimental replicates can be found in Supplementary Table 6.

## Data availability

Source data for all figures presented in the paper and Supplementary Information are available at https://doi.org/10.6084/m9.figshare.23608155.

## Code availability

Custom software is available on the Whitley lab GitHub page: https://github.com/WhitleyLab/Vercini_spt_analysis

