## Supplementary Information for "A one-track model for spatiotemporal coordination of *Bacillus subtilis* septal cell wall synthesis"

**SUPPLEMENTARY FIGURES**

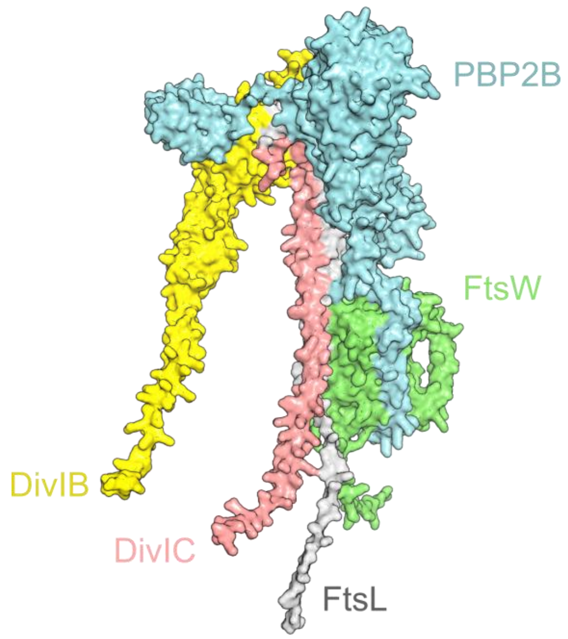

**Supplementary Figure 1. Model of the *B. subtilis* divisome synthesis complex.** Model of the pentameric *B. subtilis* divisome complex consisting of five proteins (PBP2B, cyan; FtsW, green; FtsL, grey; DivIC, pink; DivIB, yellow). Modelling details are described in Methods.

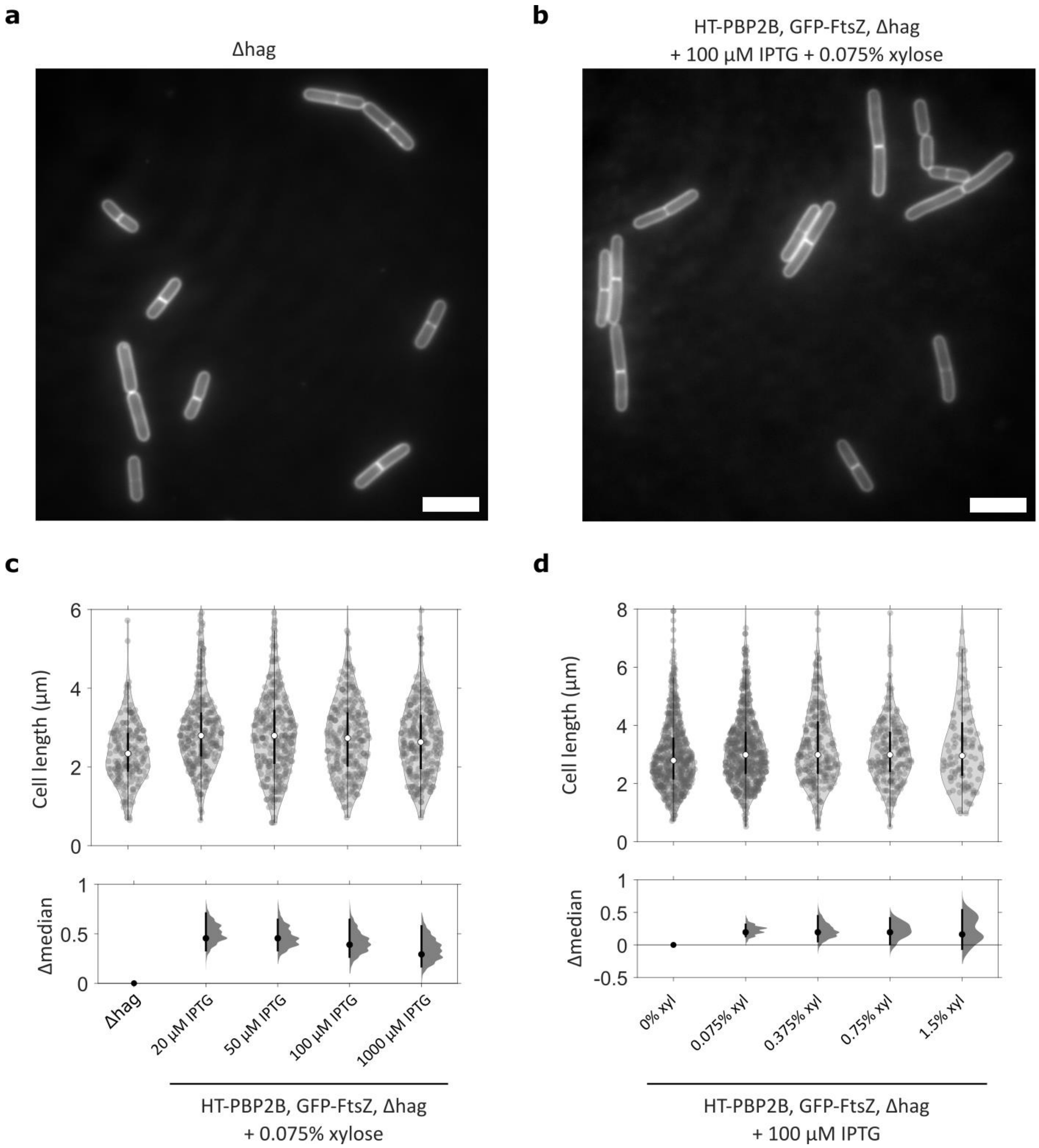

**Supplementary Figure 2. Effect of HT-PBP2B and GFP-FtsZ induction levels on cell morphology.** Lengths of HT-PBP2B GFP-FtsZ Δhag cells (strain SH147) were measured by microscopy of Nile Red-stained cells and compared to those of Δhag cells (strain SH211) (Supplementary Table 1). Both strains were grown in PHMM at 30°C. **(a)** Micrograph of *Δ*hag cells. **(b)** Micrograph of HT-PBP2B, GFP-FtsZ, Δhag cells with 100 µM IPTG and 0.075% xylose (standard experimental conditions). **(c)** HT-PBP2B expression was induced at the level specified by the [IPTG] shown, while GFP-FtsZ levels were held constant (0.075% xylose induction). **(d)** GFP-FtsZ expression was induced at the level specified by the [xyl] concentration shown, while the HT-PBP2B levels were held constant (100 µM IPTG induction). Violin plots: white circles, median; thick black lines, interquartile range; thin black lines, 1.5x interquartile range. DABEST plots: black circle, median difference between indicated conditions; black lines, 95% confidence interval of median difference. Scale bars: 5 µm. Sample sizes are listed in Supplementary Table 6.

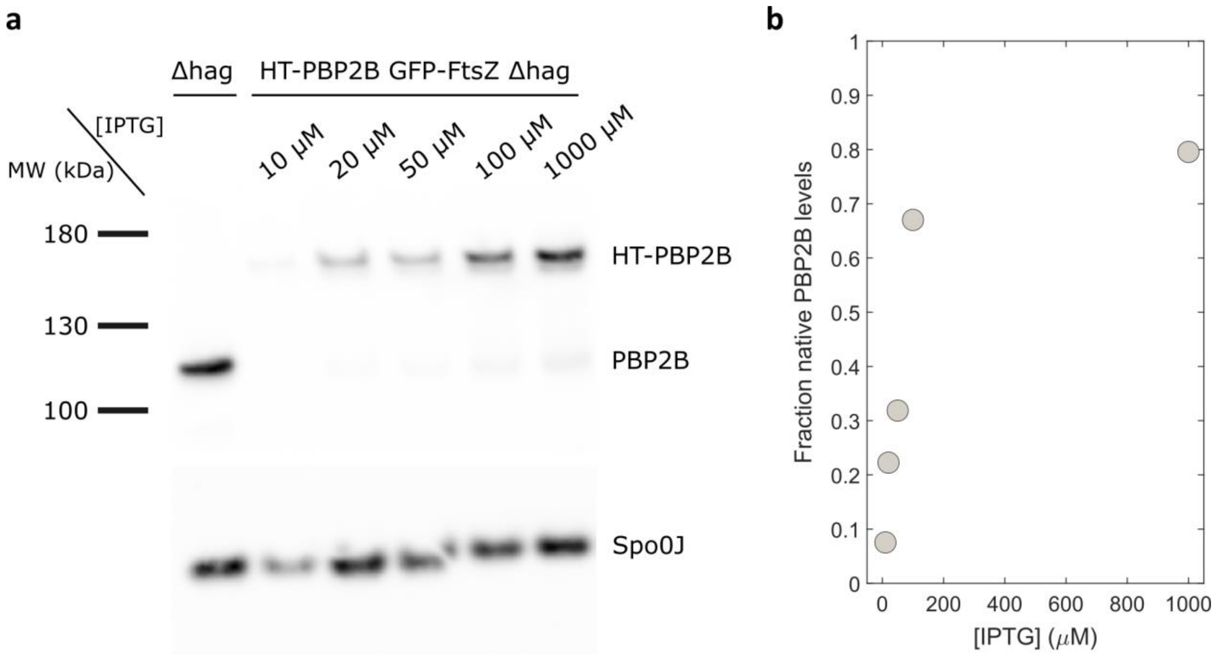

**Supplementary Figure 3. Quantification of PBP2B levels across [IPTG]. (a)** Western blot of HT-PBP2B GFP-FtsZ Δhag (strain SH147; Supplementary Table 1) and Δhag (strain SH211). Cultures were grown in rich media at 37°C with varying [IPTG] to induce HT-PBP2B expression. 5 µg of total protein from lysate was blotted. Spo0J was used as a loading control on a separate blot. The membranes were incubated with polyclonal antibodies specific for PBP2B or Spo0J. Bands were visualised by a chemiluminescence system. **(b)** Protein expression levels of HT-PBP2B across [IPTG] based on the normalized integrated intensities of bands in the Western blot. Uncropped images are provided as a Source Data file.

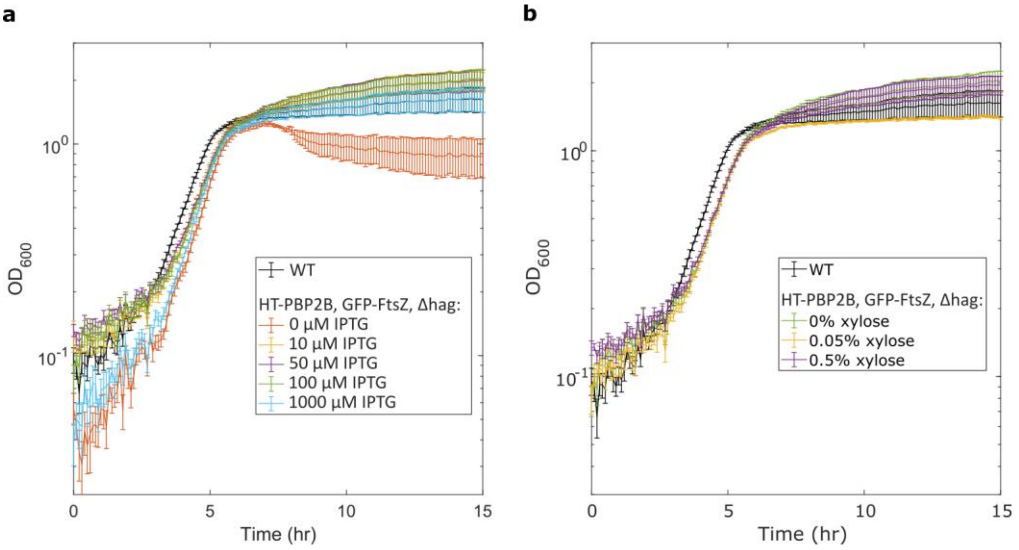

**Supplementary Figure 4. Effect of HT-PBP2B and GFP-FtsZ induction on growth in liquid culture.** Growth of HT-PBP2B GFP-FtsZ Δhag cultures (strain SH147) was monitored for 15 hours using a FLUOStar OPTIMA plate reader (BMG Labtech) and compared to that of wild-type (PY79) (Supplementary Table 1). Both strains were grown in LB at 30°C. Mean values ± SD of triplicate repeats are plotted. **(a)** HT-PBP2B expression was induced at the level specified by the [IPTG] shown, while GFP-FtsZ was not induced (0% xylose). **(b)** GFP-FtsZ expression was induced at the level specified by the [xylose] shown, while HT-PBP2B expression was induced with a constant 100 µM IPTG.

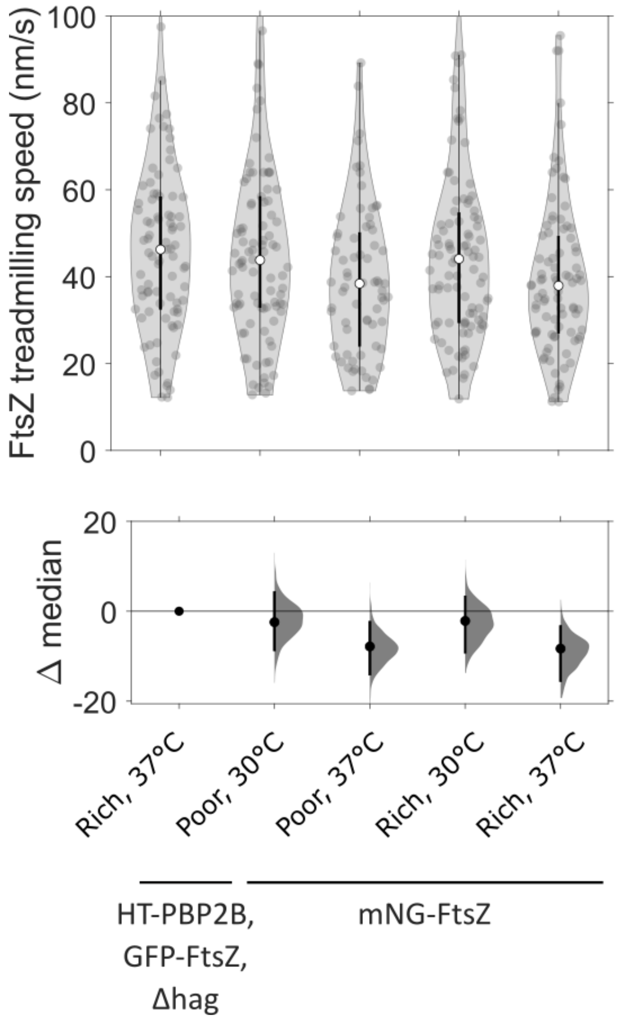

**Supplementary Figure 5. Speed of FtsZ treadmilling in HT-PBP2B, GFP-FtsZ, Δhag strain compared to previous measurements.** FtsZ treadmilling speeds were measured by TIRF microscopy using a HT-PBP2B, GFP-FtsZ, Δhag strain (strain SH147; Supplementary Table 1) and compared to treadmilling speeds in a strain expressing mNeonGreen-FtsZ (mNG-FtsZ) from an IPTG-inducible promoter (strain bWM4; Supplementary Table 1) measured across different growth conditions (poor vs. rich media, 30°C vs. 37 °C). HT-PBP2B, GFP-FtsZ, Δhag cultures were grown in rich media (PHMM) at 37°C with 0.075% xylose to induce a low level of GFP-FtsZ expression. Treadmilling speeds were determined by manually tracing filament trajectories on kymographs, as done previously[2]. Data for all mNG-FtsZ conditions are taken from Whitley et al[2]. Violin plots: white circles, median; thick black lines, interquartile range; thin black lines, 1.5x interquartile range. DABEST plots: black circle, median difference between indicated conditions; black lines, 95% confidence interval of median difference. Sample sizes are listed in Supplementary Table 6.

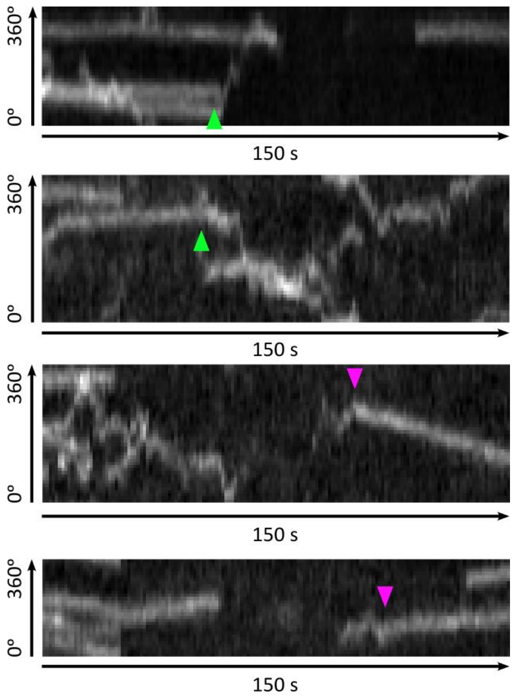

**Supplementary Figure 6. Transitions to and from the diffusive state.** Four example radial kymographs showing HT-PBP2B molecules transitioning from either an immobile or processive state to a diffusive state (green arrowheads), or from a diffusive state to a processive state (magenta arrowheads).

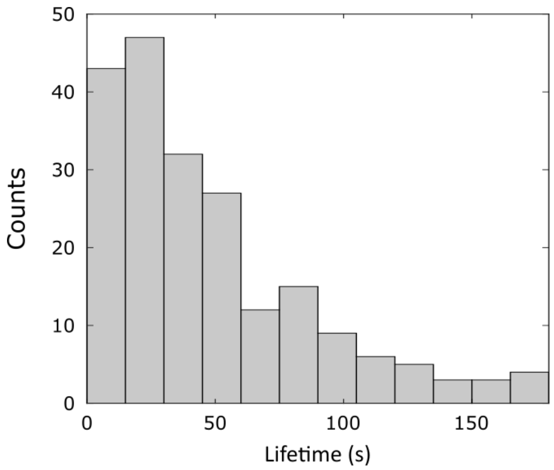

**Supplementary Figure 7. Lifetime of immobile HT-PBP2B tracks.** Histogram of lifetimes for immobile track segments from HT-PBP2B GFP-FtsZ Δhag cells (strain SH147) grown in PHMM at 30°C. Here, immobile is defined as a track segment with speed less than 4 nm/s. Sample sizes are listed in Supplementary Table 6.

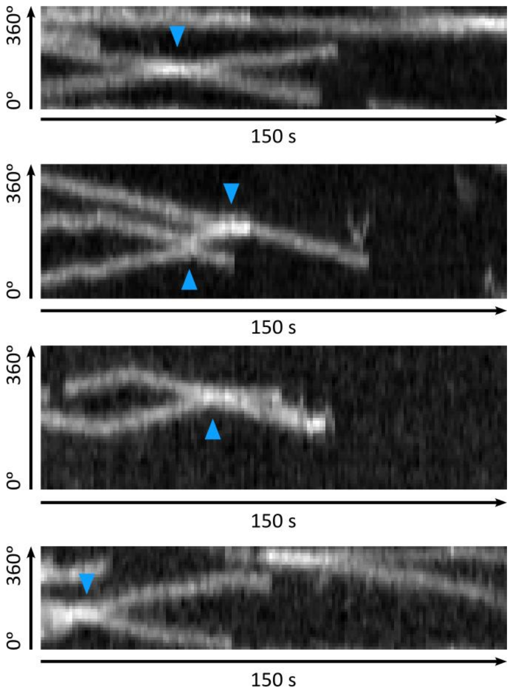

**Supplementary Figure 8. Track crossing events.** Four examples of radial kymographs of HT-PBP2B showing processive tracks crossing over one another (crossing events shown with blue arrowheads).

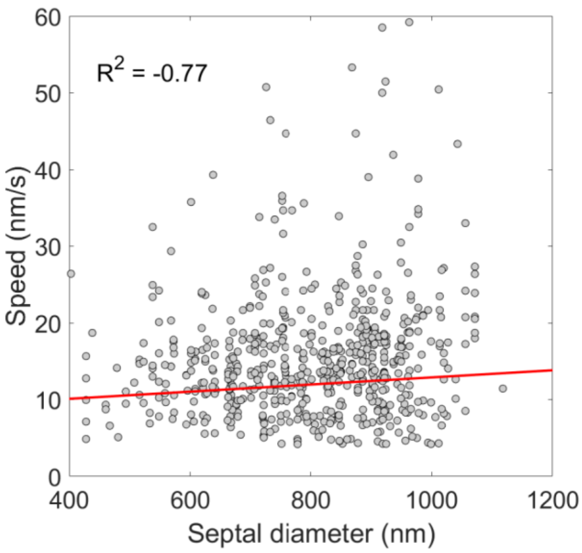

**Supplementary Figure 9. Correlation between HT-PBP2B speed and septal diameter.** Scatter plots showing the speeds of processive HT-PBP2B track segments (speeds between 4 and 60 nm/s) and the associated diameters of the septa around which they were moving under our standard conditions (rich media, 30°C).

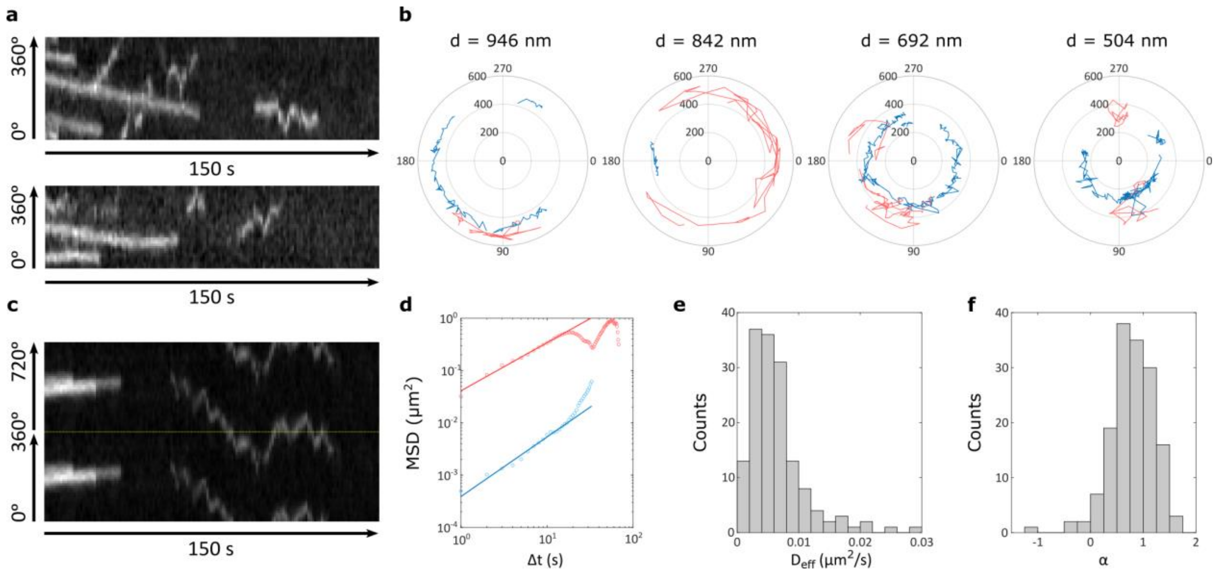

**Supplementary Figure 10. Diffusive motion of HT-PBP2B. (a)** Two example radial kymographs of HT-PBP2B (rich media, 30°C) showing both processive and diffusive tracks. Diffusive tracks appear as short back-and-forth lines. **(b)** Examples of single-molecule tracks of HT-PBP2B across four septal diameters, plotted in polar coordinates. Tracks were produced using TrackMate[1] with 0.5 µm linking distance and 0 frame gaps, then plotted in polar coordinates using bespoke MATLAB code. All processive and immobile tracks are shown in blue, while diffusive tracks are shown in red. **(c)** Example radial kymograph of HT-PBP2B (rich media, 30°C) showing a processive track and a long-lived diffusive track. As the diffusive track covered most of the cell circumference, two revolutions around the cell (0°-360° and 360°-720°) are plotted top and bottom, separated by a dotted yellow line. **(d)** Example mean-squared displacement (MSD) vs. time step plot for the kymograph shown in panel c. Blue circles: MSDs from processive track. Red circles: MSDs from diffusive track. Blue line: fit to MSDs from processive track. Red line: fit to MSDs from diffusive track (fit details in Methods). **(e)** Histogram of effective diffusion coefficients from MSD analyses. **(f)** Histogram of anomalous diffusion exponents $\alpha$ from MSD analyses. Sample sizes are listed in Supplementary Table 6.

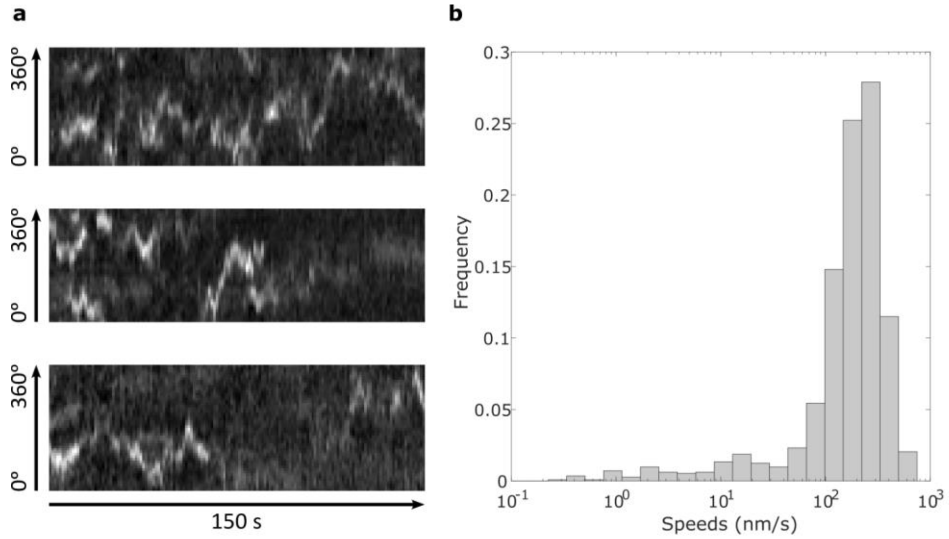

**Supplementary Figure 11. Motion of HT-PBP2B molecules outside the septal ring area. (a)** Example radial kymographs of HT-PBP2B (rich media, 30°C) showing motion outside the septal ring area (*i.e.* at a random section of the cell sidewall). **(b)** Histogram of HT-PBP2B speeds for molecules observed outside the septal ring area plotted on logarithmic x axis, measured from linear segments on kymographs.

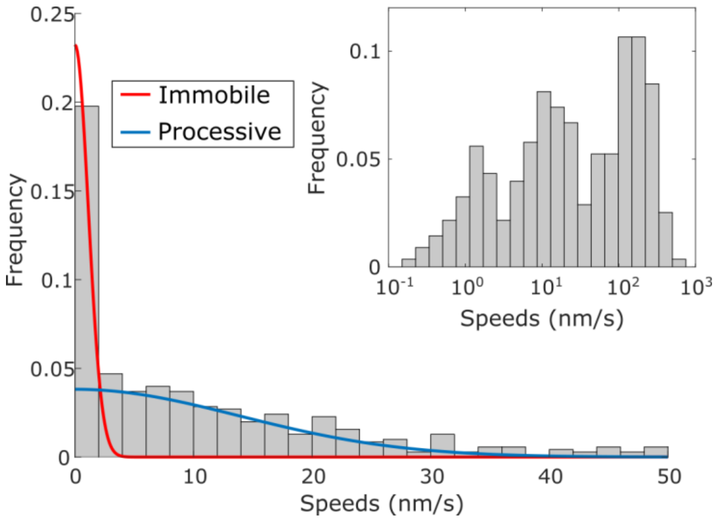

**Supplementary Figure 12. Speeds of HT-PBP2B molecules with high inducer concentration.** Histogram of HT-PBP2B speeds in HT-PBP2B GFP-FtsZ Δhag cells (strain SH147; Supplementary Table 1) grown in rich media with 1 mM IPTG and 0.075% xylose at 30°C with 250-500 pM JFX554 HaloTag ligand and imaged using VerCINI. Red and blue lines show fits to the data. *Inset*: Histogram of speeds plotted on logarithmic x axis, showing three populations.

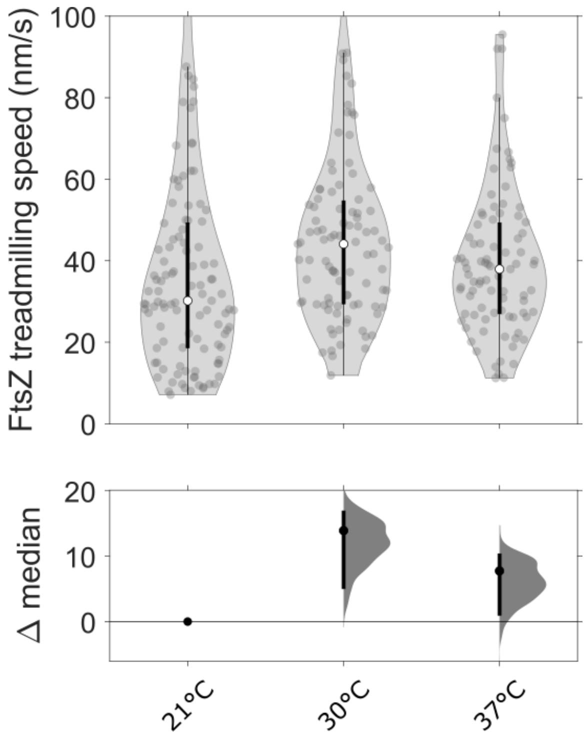

**Supplementary Figure 13. Effect of temperature on FtsZ treadmilling speed.** FtsZ treadmilling speeds were measured by TIRF microscopy using a strain expressing mNeonGreen-FtsZ from an IPTG-inducible promoter (strain bWM4; Supplementary Table 1) grown in rich media at 21°C and compared to previous measurements of the same strain in the same media under varying temperatures. Cultures were incubated with 25 μM IPTG 1 hr prior to imaging to induce mNeonGreen-FtsZ expression. Treadmilling speeds were determined by manually tracing filament trajectories on kymographs, as done previously[2]. Data for 30°C and 37°C are taken from Whitley et al. 2021[2]. Violin plots: white circles, median; thick black lines, interquartile range; thin black lines, 1.5x interquartile range. DABEST plots: black circle, median difference between indicated conditions; black lines, 95% confidence interval of median difference. Sample sizes are listed in Supplementary Table 6.

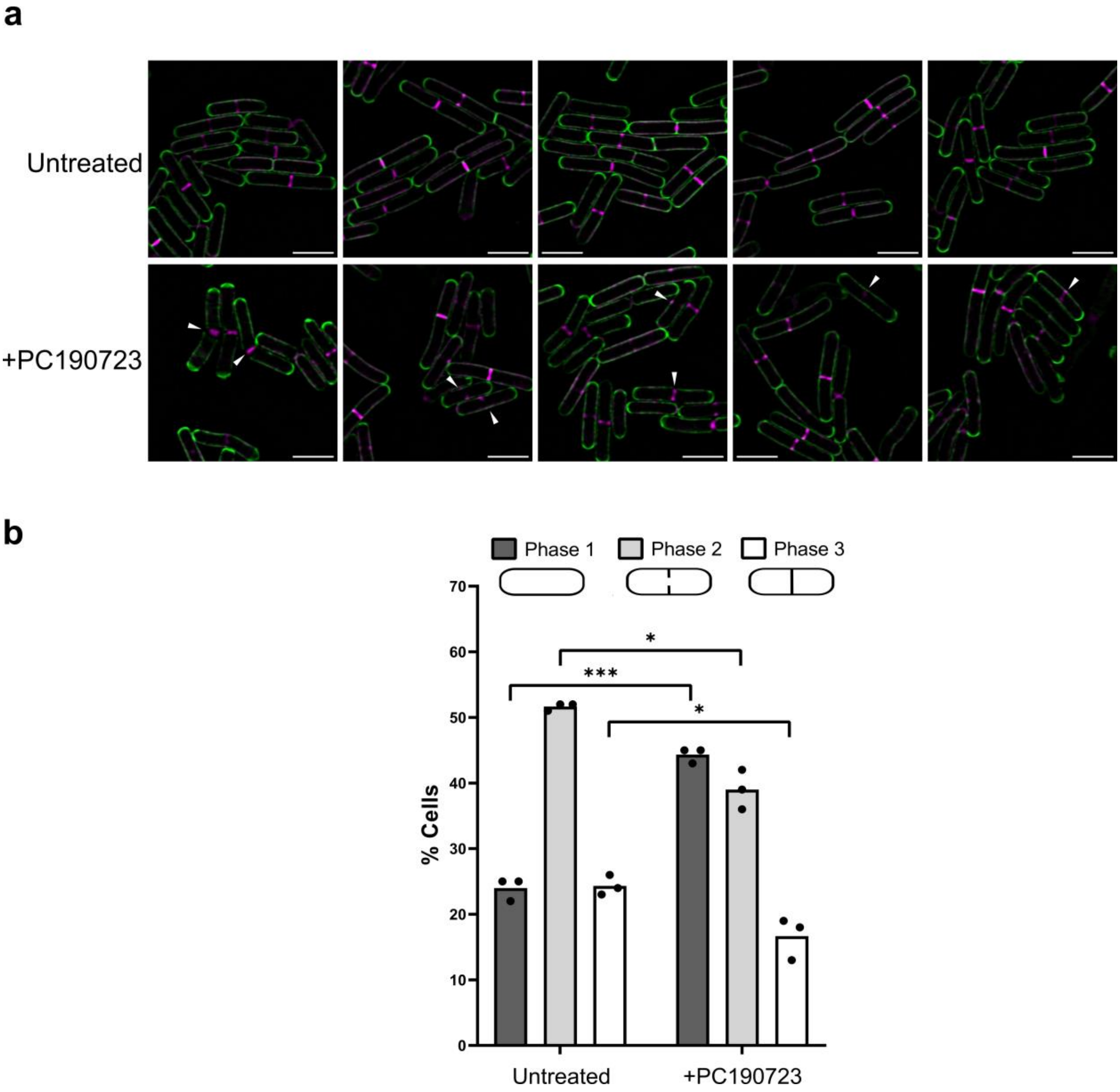

**Supplementary Figure 14. FtsZ treadmilling is not required for septal PG synthesis.** Cells (strain PY79; Supplementary Table 1) were stained with the fluorescent D-amino acid BADA for 90 min, then with TADA for 10 min. To arrest treadmilling by FtsZ, cells were treated with 14 µM PC190723 (5×MIC) for 10 min during the TADA staining step. **(a)** Panels show representative images of stained cells. White arrows show septal aberrations. Scale bars: 3 µm. **(b)** Quantification of the percentage of cells in each division phase in cells stained with BADA and TADA with or without PC190723. At least 100 cells were counted per repeat. Sample sizes are listed in Supplementary Table 6. The histograms show the means, and *p*-values are a result of unpaired *t*-tests with Welch's correction (two-tailed). *: $p < 0.05$, ***: $p < 0.001$.

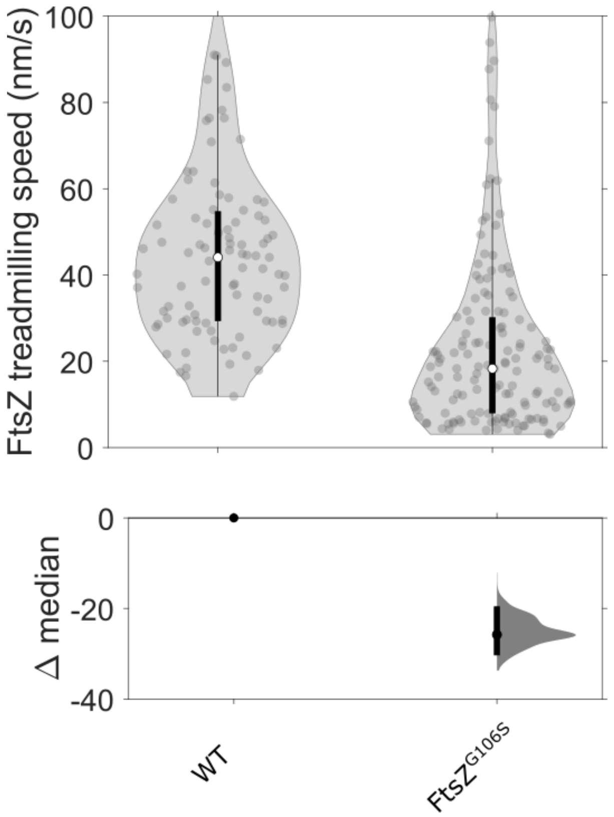

**Supplementary Figure 15. Effect of FtsZ$^{G106S}$ mutation on treadmilling speed.** FtsZ treadmilling speeds were measured for cells expressing FtsZ$^{G106S}$ (strain SH203; Supplementary Table 1) in rich media at 30°C and compared to treadmilling speeds in a strain expressing mNeonGreen-FtsZ from an IPTG-inducible promoter (strain bWM4) measured under the same conditions. FtsZ$^{G106S}$ cells were incubated with 0.075% xylose to induce GFP-FtsZ expression. Treadmilling speeds were determined by manually tracing filament trajectories on kymographs, as done previously[2]. Data for WT are taken from Whitley et al. 2021[2]. Violin plots: white circles, median; thick black lines, interquartile range; thin black lines, 1.5x interquartile range. DABEST plots: black circle, median difference between indicated conditions; black lines, 95% confidence interval of median difference. Sample sizes are listed in Supplementary Table 6.

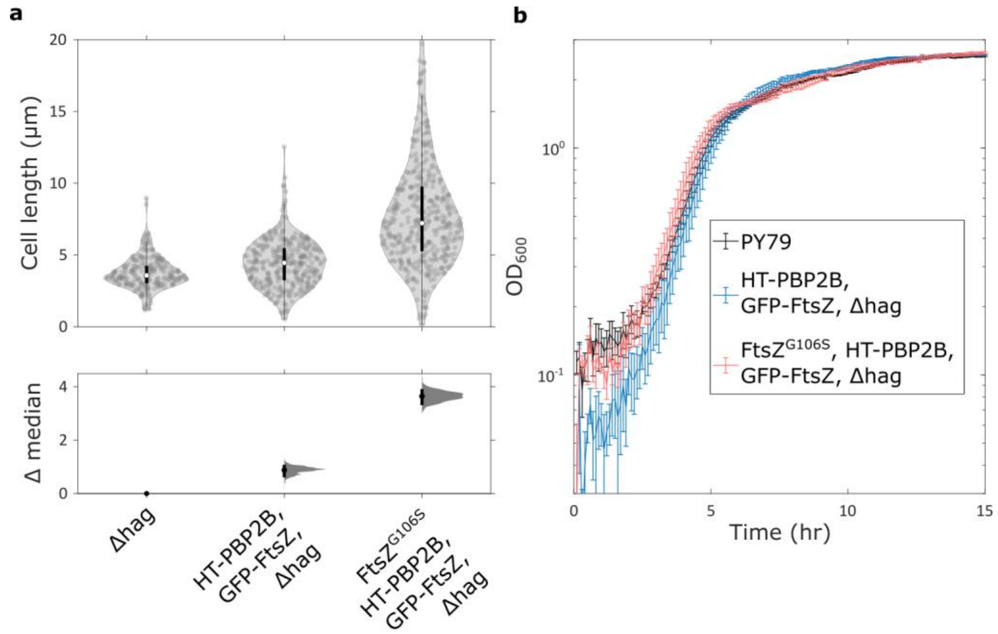

**Supplementary Figure 16. Effect of FtsZ$^{G106S}$ mutation on cell lengths and growth in liquid culture. (a)** Lengths of FtsZ$^{G106S}$ HT-PBP2B GFP-FtsZ Δhag cells (strain SH203) were measured by microscopy of Nile Red-stained cells and compared to those of HT-PBP2B GFP-FtsZ Δhag and Δhag cells (strains SH147 and SH211, respectively). All strains were grown in PHMM at 30°C. HT-PBP2B expression was induced with 100 µM IPTG in both strains containing HT-PBP2B, while GFP-FtsZ was not induced in either case (0% xylose). Violin plots: white circles, median; thick black lines, interquartile range; thin black lines, 1.5x interquartile range. DABEST plots: black circle, median difference between indicated conditions; black lines, 95% confidence interval of median difference. Sample sizes are listed in Supplementary Table 6. **(b)** Growth was monitored for 15 hours using a FLUOStar OPTIMA plate reader (BMG Labtech) at 30°C. Mean values ± SD of triplicate repeats are plotted. All strains were grown in LB at 30°C. HT-PBP2B expression was induced with 100 µM IPTG in both strains containing HT-PBP2B, while GFP-FtsZ was not induced in either case (0% xylose).

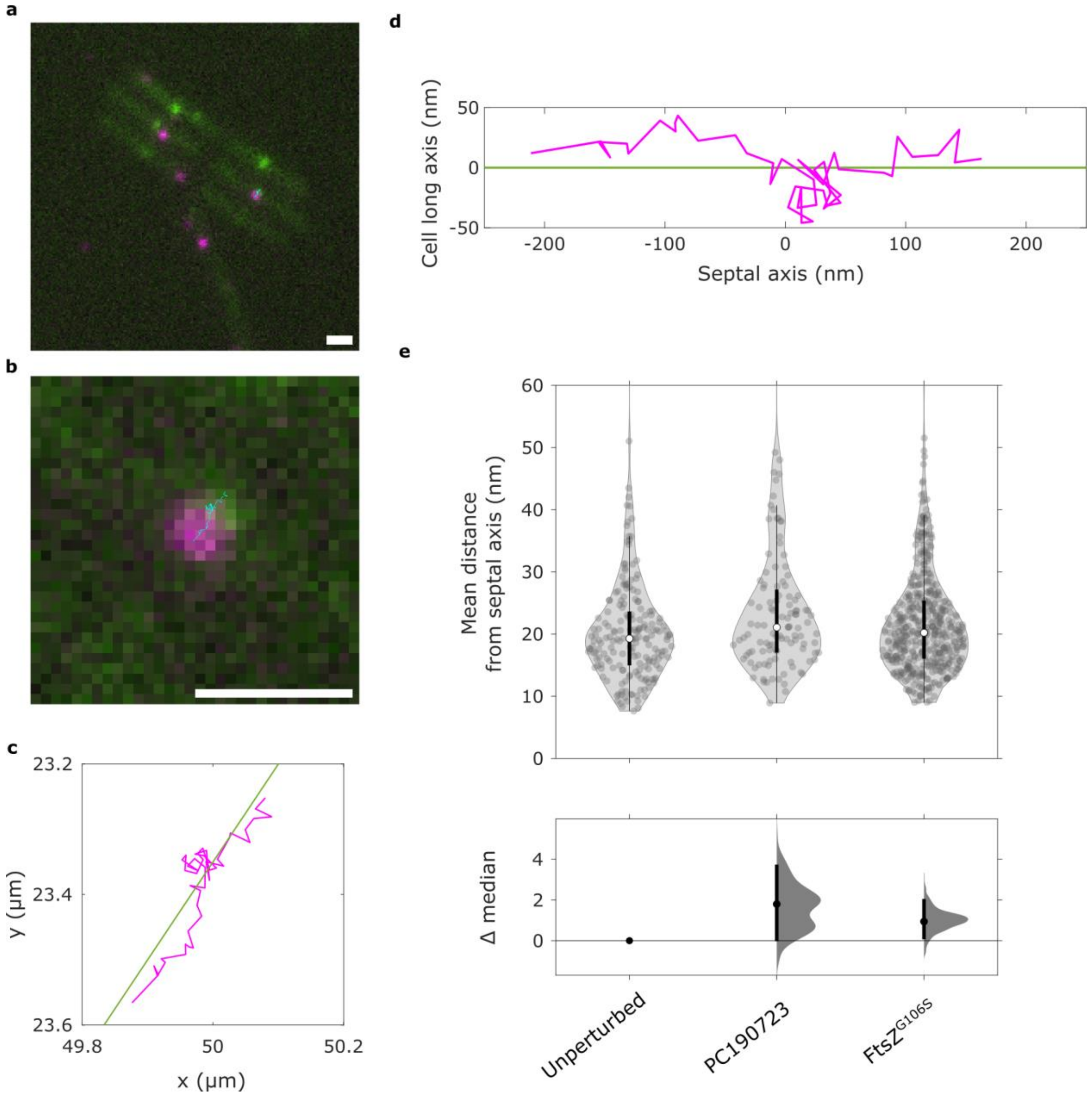

**Supplementary Figure 17. Motion of HT-PBP2B along the septal axis in horizontally-oriented cells. (a)** Example image of HT-PBP2B GFP-FtsZ Δhag strain (strain SH147; Supplementary Table 1) from a single-particle tracking video acquired with TIRF illumination (rich media, 30°C). Shown is a merge of channels for imaging GFP-FtsZ (green) and HT-PBP2B (magenta). A single-molecule track is shown in cyan. Track were produced using TrackMate[1] with 0.1 µm linking distance and 0 frame gaps. Processive molecules were selected using the filters: >9 spots in track, >90 nm track displacement, <60 nm/s median track speed. **(b)** Zoom-in of single-molecule track of HT-PBP2B from panel **a** along a division septum. **(c)** Coordinates of the example track (magenta) along with a line showing the septal axis (green). **(d)** Example track from panel **c** rotated to show coordinates along the septal axis. **(e)** Mean distances of HT-PBP2B localizations from septal axes across conditions affecting FtsZ treadmilling. Each point represents the mean distance from the septal axis from a single processive HT-PBP2B track (e.g. the track in panel **d** represents one point). Violin plots: white circles, median; thick black lines, interquartile range; thin black lines, 1.5x interquartile range. DABEST plots: black circle, median difference between indicated conditions; black lines, 95% confidence interval of median difference. Microscope acquisition parameters are listed in Supplementary Table 5. Sample sizes are listed in Supplementary Table 6. Scale bars: 1 µm.

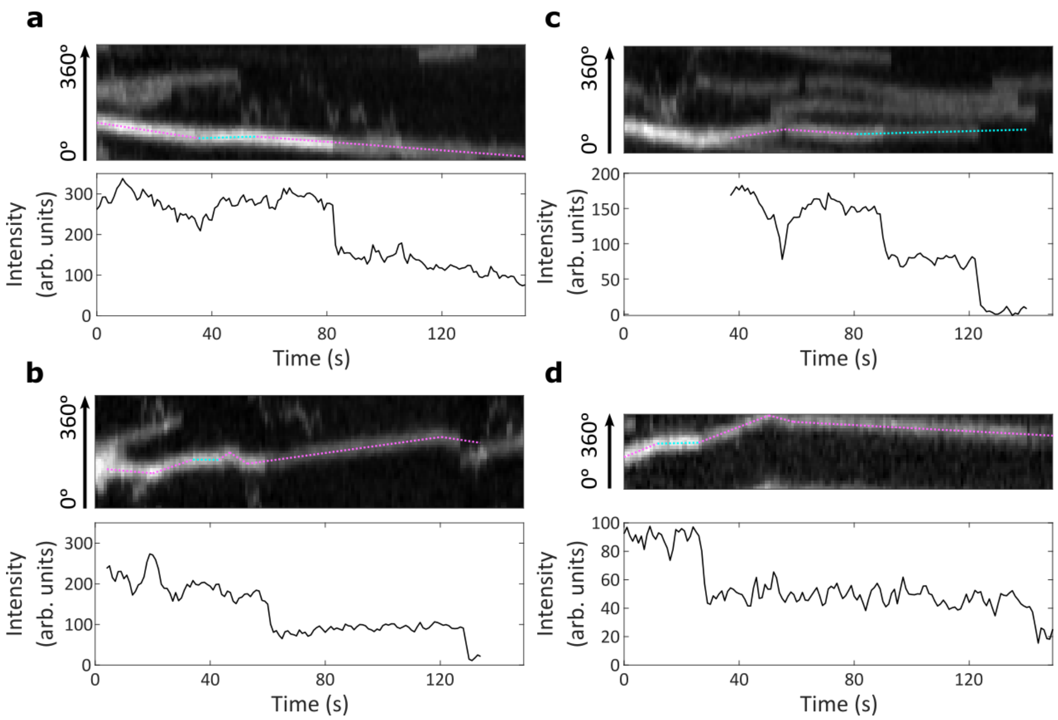

**Supplementary Figure 18. Fluorescence intensity drops during motion and after direction changes.** Four example kymographs and accompanying intensity plots are shown. *Top:* Example kymographs of HT-PBP2B motion from smVerCINI videos showing discrete drops in fluorescence intensity during motion or after direction changes. Magenta segments: processive motion. Cyan segments: immobile. *Bottom:* Intensity traces for the tracks designated by dotted cyan lines overlaid on kymographs.

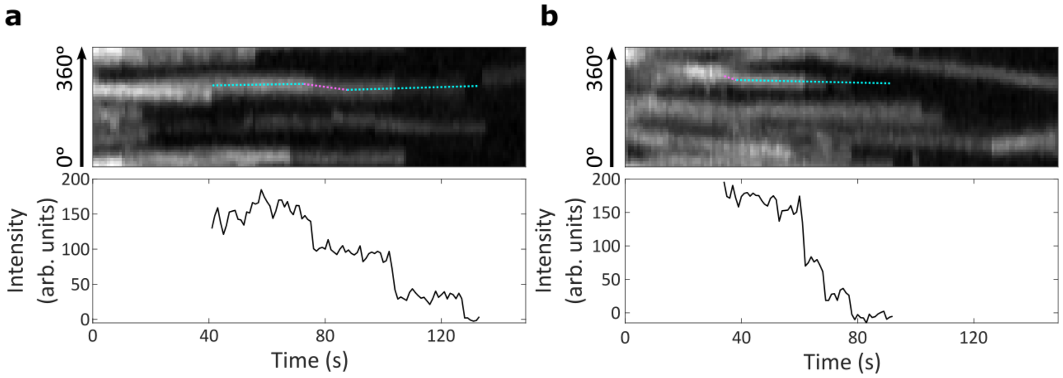

**Supplementary Figure 19. Multiple fluorescence intensity drops from single spots.** Two example kymographs and accompanying intensity plots are shown. *Top:* Example kymographs of HT-PBP2B motion from smVerCINI videos showing several discrete drops in fluorescence intensity. Magenta segments: processive motion. Cyan segments: immobile. *Bottom:* Intensity traces for the tracks designated by dotted cyan lines overlaid on kymographs.

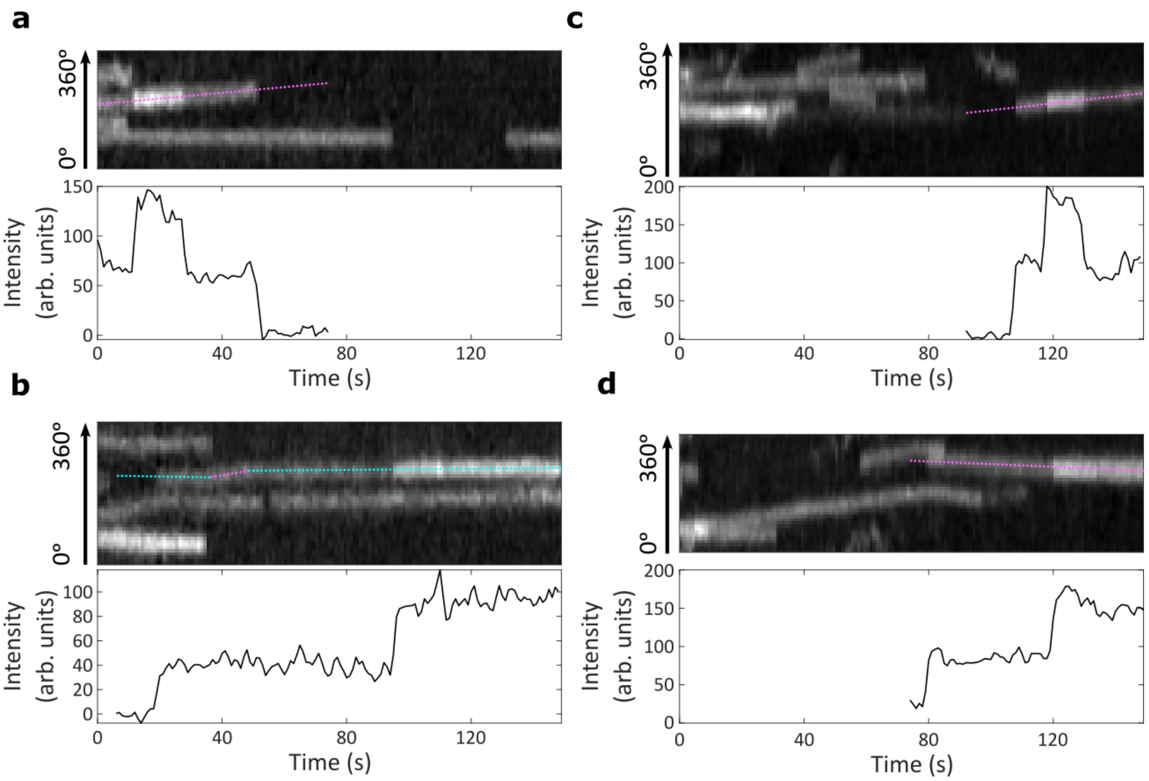

**Supplementary Figure 20. Fluorescence intensity jumps.** Four example kymographs and accompanying intensity plots are shown. *Top:* Example kymographs of HT-PBP2B motion from smVerCINI videos showing discrete jumps in fluorescence intensity. Magenta segments: processive motion. Cyan segments: immobile. *Bottom:* Intensity traces for the tracks designated by dotted cyan lines overlaid on kymographs.

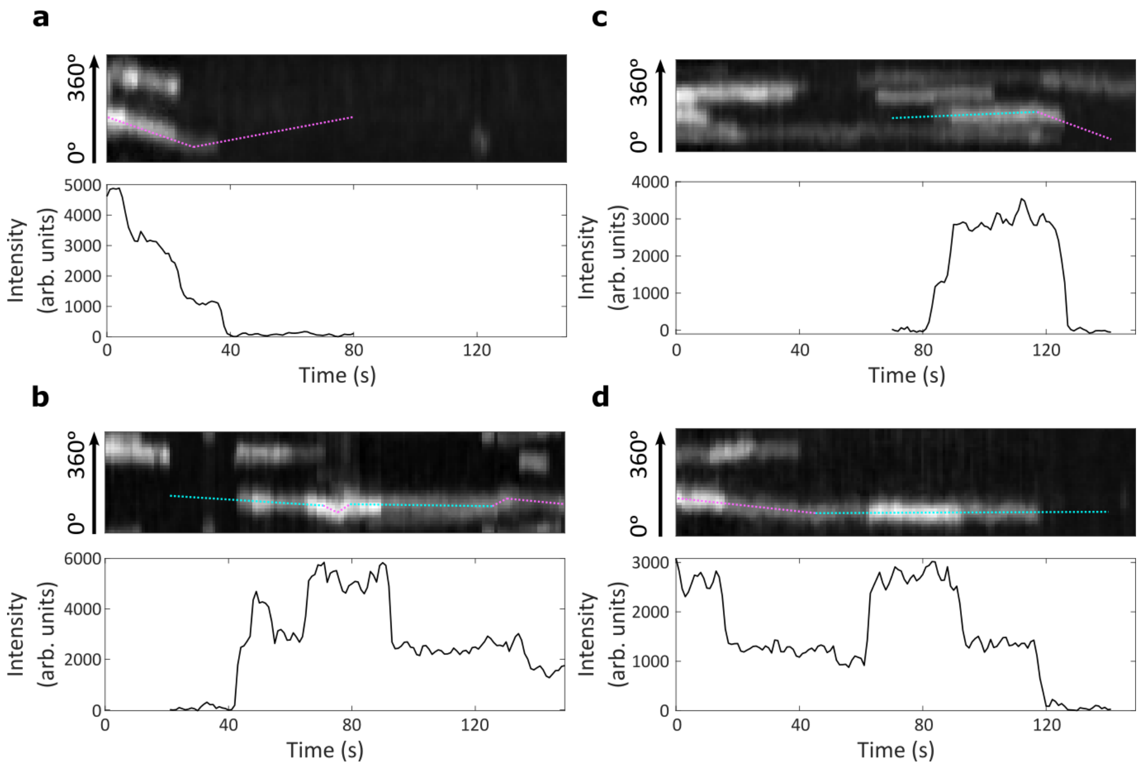

**Supplementary Figure 21. Fluorescence intensity drops and jumps in HT-FtsW tracks.** Four example kymographs and accompanying intensity plots are shown for tracks of JFX554-HaloTag-FtsW (strain bAB350; Supplementary Table 1). Cells were incubated with 5 nM JFX554 HaloTag ligand for 15 min prior to imaging. *Top:* Example kymographs of HT-FtsW motion from smVerCINI videos showing discrete drops or jumps in fluorescence intensity. Magenta segments: processive motion. Cyan segments: immobile. *Bottom:* Intensity traces for the tracks designated by dotted cyan lines overlaid on kymographs.

**SUPPLEMENTARY VIDEO LEGENDS**

**Supplementary Videos 1 and 2. Example videos of single-molecule tracks of HT-PBP2B in vertically-trapped cells.** Each video shows a single division septum of a HT-PBP2B GFP-FtsZ Δhag cell (strain SH147; Supplementary Table 1) in rich media 30°C with 100 µM IPTG and 0.075% xylose. HT-PBP2B is labelled sub-stoichiometrically with JFX554 HaloTag ligand (250 pM) so that single molecules can be easily observed. Videos show acquisition with a 561 nm laser operating in ring-HiLO illumination mode with 500 ms camera exposure time at 1 s frame intervals (Supplementary Table 5). Supplementary Video 1 corresponds to the top kymograph in Figure 1c of the main text, while Video 2 corresponds to the bottom kymograph, as well as Figure 4a. Scale bars: 1 µm.

**Supplementary Videos 3 and 4. Example videos of single-molecule tracks of HT-PBP2B in vertically-trapped cells with Penicillin G treatment.** Each video shows a single division septum of a HT-PBP2B GFP-FtsZ Δhag cell (strain SH147; Supplementary Table 1) in rich media 30°C with 100 µM IPTG and 0.075% xylose with excess penicillin G (20 µg/mL). HT-PBP2B is labelled sub-stoichiometrically with JFX554 HaloTag ligand (250 pM) so that single molecules can be easily observed. Videos show acquisition with a 561 nm laser operating in ring-HiLO illumination mode with 500 ms camera exposure time at 1 s frame intervals (Supplementary Table 5). Video 3 corresponds to the top kymograph in Figure 1d of the main text, while Video 4 corresponds to the bottom kymograph. Scale bars: 1 µm.

**Supplementary Videos 5 and 6. Example videos of single-molecule tracks of HT-PBP2B in vertically-trapped cells with fosfomycin treatment.** Each video shows a single division septum of a HT-PBP2B GFP-FtsZ Δhag cell (strain SH147; Supplementary Table 1) in rich media 30°C with 100 µM IPTG and 0.075% xylose with excess fosfomycin (500 µg/mL). HT-PBP2B is labelled sub-stoichiometrically with JFX554 HaloTag ligand (250 pM) so that single molecules can be easily observed. Videos show acquisition with a 561 nm laser operating in HiLO illumination mode with 500 ms camera exposure time at 1 s frame intervals (Supplementary Table 5). Video 5 corresponds to the top kymograph in Figure 1e of the main text, while Video 6 corresponds to the bottom kymograph. Scale bars: 1 µm.

**Supplementary Videos 7 and 8. Example videos of single-molecule tracks of HT-PBP2B in vertically-trapped cells with PC190723 treatment.** Each video shows a single division septum of a HT-PBP2B GFP-FtsZ Δhag cell (strain SH147; Supplementary Table 1) in rich media 30°C with 100 µM IPTG and 0.075% xylose with excess PC190723 (10 µM). HT-PBP2B is labelled sub-stoichiometrically with JFX554 HaloTag ligand (250 pM) so that single molecules can be easily observed. Videos show acquisition with a 561 nm laser operating in ring-HiLO illumination mode with 500 ms camera exposure time at 1 s frame intervals (Supplementary Table 5). Video 7 corresponds to the top kymograph in Figure 3a of the main text, while Video 8 corresponds to the bottom kymograph. Scale bars: 1 µm.

**Supplementary Videos 9 and 10. Example videos of single-molecule tracks of HT-PBP2B in vertically-trapped cells expressing FtsZ$^{G106S}$.** Each video shows a single division septum of an FtsZ$^{G106S}$ HT-PBP2B GFP-FtsZ Δhag cell (strain SH203; Supplementary Table 1) in rich media 30°C with 100 µM IPTG and 0.075% xylose. HT-PBP2B is labelled sub-stoichiometrically with JFX554 HaloTag ligand (250 pM) so that single molecules can be easily observed. Videos show acquisition with a 561 nm laser operating in HiLO illumination mode with 500 ms camera exposure time at 1 s frame intervals (Supplementary Table 5). Video 9 corresponds to the top kymograph in Figure 3b of the main text, while Video 10 corresponds to the bottom kymograph. Scale bars: 1 µm.

**Supplementary Videos 11-13. Example videos of single-molecule tracks of HT-PBP2B in vertically-trapped cells showing fluorescence intensity drops.** Each video shows a single division septum of a HT-PBP2B GFP-FtsZ Δhag cell (strain SH147; Supplementary Table 1) at 30°C with 100 µM IPTG and

0.075% xylose. Videos show acquisition with a 561 nm laser operating in ring-HiLO illumination mode with 500 ms camera exposure time at 1 s frame intervals (Supplementary Table 5). Supplementary Video 11 (rich media, 250 pM JFX554 HaloTag ligand) corresponds to the kymograph in Figure 4b of the main text. Videos 12 and 13 (minimal media, 100 pM JFX554 HaloTag ligand) correspond to the kymographs in Figure 4c and Figure 4d respectively. Scale bars: 1 µm.

**Supplementary Videos 14-16: Example videos of single-molecule tracks of HT-PBP2B in horizontally-oriented cells showing motion along the septal axis.** Each video shows a field of view covering several cells in rich media at 30°C with 100 µM IPTG and 0.075% xylose. In each case, HT-PBP2B is labelled sub-stoichiometrically with JFX554 HaloTag ligand 250 pM) so that single molecules can be easily observed. Videos show a still image with a 488 nm laser operating in TIRF illumination mode with 1-5 s exposure time (green), merged with a video taken with a 561 nm laser operating in TIRF illumination mode with 500 ms exposure time and 1 s frame interval (magenta) (Supplementary Table 5). Molecules were identified and tracks established using TrackMate. Tracks for molecules passing thresholds (described in legend to Supplementary Figure 17) are depicted in cyan. Video 14: unperturbed HT-PBP2B GFP-FtsZ Δhag (strain SH147; Supplementary Table 1) cells, corresponding to the images and analysis shown in Supplementary Figure 17. Video 15: PC190723-treated HT-PBP2B GFP-FtsZ Δhag cells (10 µM PC190723). Video 16: $\text{FtsZ}^{\text{G106S}}$ HT-PBP2B GFP-FtsZ Δhag cells. Scale bars: 1 µm

### SUPPLEMENTARY TABLES

**Supplementary Table 1: Strains used in this study.**

| Strain name | Genotype/Description[a] | Source |
|---|---|---|
| PY79 | Prototroph | [3] |
| SH211 | PY79 Δ*hag* | [2] |
| 2020 | 168 *trpC2*, *amyE*::*spc*-$P_{xyl}$-*gfp-ftsZ* | [4] |
| SH142 | PY79 Δ*hag amyE*::*spc*-$P_{xyl}$-*gfp-ftsZ* | This work |
| bGS31 | PY79 *pbpB*::*erm*-$P_{hyperspank}$-*HaloTag-15aa-pbpB, ftsZ*::*mNeonGreen-15aa-ftsZ* multicopy | [5] |
| SH147 | PY79 Δ*hag pbpB*::*erm*-$P_{hyperspank}$-*HaloTag-15aa-pbpB*, *amyE*::*spc*-$P_{xyl}$-*gfp-ftsZ* | This work |
| Z-G106S | PY79 *ftsZΩftsZ(G106S) (tet)* | Gift from Ethan Garner (Harvard) |
| SH203 | PY79 Δ*hag pbpB*::*erm*-$P_{hyperspank}$-*HaloTag-15aa-pbpB*, *ftsZΩftsZ(G106S) (tet), amyE*::*spc*-$P_{xyl}$-*gfp-ftsZ* | This work |
| bWM4 | PY79 *amyE::erm*-$P_{hyperspank}$-*ftsA-mNeonGreen-15aa-ftsZ* | [5] |
| bAB350 | PY79 *ftsW::erm*-$P_{xyl}$-*HaloTag-15aa-ftsW* | [6] |

[a]Drug resistance cassettes: erm, erythromycin resistance; spc, spectinomycin resistance; tet, tetracycline resistance

**Supplementary Table 2: Probabilities of transition between states and transition rates for each HT-PBP2B motion state under each experimental condition.** Rate constants were calculated using the formula in Supplementary Note 1. Errors are SEM.

| Condition | Transition | Probability | Rate constant ($s^{-1}$) |
|---|---|---|---|
| Poor media 30°C | Immobile -> Signal loss | 0.88 ± 0.08 | 0.015 ± 0.002 |
| | Immobile -> Processive | 0.10 ± 0.02 | 0.0018 ± 0.0004 |

| | | | |
|---|---|---|---|
| | Immobile -> Diffusive | 0.012 ± 0.007 | 0.0002 ± 0.0001 |
| | Processive -> Signal loss | 0.55 ± 0.06 | 0.014 ± 0.002 |
| | Processive -> Processive (direction change) | 0.17 ± 0.03 | 0.0043 ± 0.0007 |
| | Processive -> Immobile | 0.25 ± 0.03 | 0.0061 ± 0.0009 |
| | Processive -> Diffusive | 0.02 ± 0.01 | 0.0006 ± 0.0002 |
| | Processive -> Processive (speed change) | 0 | 0 |
| Poor media 37°C | Immobile -> Signal loss | 0.85 ± 0.07 | 0.013 ± 0.001 |
| | Immobile -> Processive | 0.12 ± 0.02 | 0.0019 ± 0.0003 |
| | Immobile -> Diffusive | 0.03 ± 0.01 | 0.0006 ± 0.0002 |
| | Processive -> Signal loss | 0.42 ± 0.03 | 0.014 ± 0.001 |
| | Processive -> Processive (direction change) | 0.32 ± 0.03 | 0.011 ± 0.001 |
| | Processive -> Immobile | 0.23 ± 0.02 | 0.0074 ± 0.0008 |
| | Processive -> Diffusive | 0.021 ± 0.006 | 0.0007 ± 0.0002 |
| | Processive -> Processive (speed change) | 0.008 ± 0.004 | 0.0002 ± 0.0001 |
| Rich media 30°C | Immobile -> Signal loss | 0.89 ± 0.08 | 0.020 ± 0.002 |
| | Immobile -> Processive | 0.07 ± 0.02 | 0.0015 ± 0.0004 |
| | Immobile -> Diffusive | 0.04 ± 0.01 | 0.0009 ± 0.0003 |
| | Processive -> Signal loss | 0.54 ± 0.04 | 0.019 ± 0.002 |
| | Processive -> Processive (direction change) | 0.29 ± 0.03 | 0.010 ± 0.001 |
| | Processive -> Immobile | 0.12 ± 0.02 | 0.0042 ± 0.0006 |
| | Processive -> Diffusive | 0.05 ± 0.01 | 0.0017 ± 0.0003 |
| | Processive -> Processive (speed change) | 0.005 ± 0.003 | 0.0002 ± 0.0001 |
| Rich media 37°C | Immobile -> Signal loss | 0.86 ± 0.07 | 0.023 ± 0.002 |
| | Immobile -> Processive | 0.10 ± 0.02 | 0.0025 ± 0.0005 |
| | Immobile -> Diffusive | 0.04 ± 0.01 | 0.0010 ± 0.0003 |
| | Processive -> Signal loss | 0.49 ± 0.03 | 0.024 ± 0.002 |

| | | | |
|---|---|---|---|
| | Processive -> Processive (direction change) | 0.36 ± 0.03 | 0.017 ± 0.001 |
| | Processive -> Immobile | 0.10 ± 0.01 | 0.0050 ± 0.0006 |
| | Processive -> Diffusive | 0.033 ± 0.007 | 0.0016 ± 0.0003 |
| | Processive -> Processive (speed change) | 0.007 ± 0.003 | 0.0003 ± 0.0001 |
| Rich media 30°C + penicillin G | Immobile -> Signal loss | 0.97 ± 0.05 | 0.025 ± 0.002 |
| | Immobile -> Processive | 0.009 ± 0.004 | 0.0002 ± 0.0001 |
| | Immobile -> Diffusive | 0.022 ± 0.006 | 0.0006 ± 0.0002 |
| | Processive -> Signal loss | 0.6 ± 0.1 | 0.06 ± 0.01 |
| | Processive -> Processive (direction change) | 0.14 ± 0.05 | 0.015 ± 0.005 |
| | Processive -> Immobile | 0.10 ± 0.04 | 0.010 ± 0.004 |
| | Processive -> Diffusive | 0.20 ± 0.06 | 0.021 ± 0.007 |
| | Processive -> Processive (speed change) | 0 | 0 |
| Rich media 30°C + fosfomycin | Immobile -> Signal loss | 0.99 ± 0.06 | 0.033 ± 0.003 |
| | Immobile -> Processive | 0.010 ± 0.005 | 0.0003 ± 0.0002 |
| | Immobile -> Diffusive | 0 | 0 |
| | Processive -> Signal loss | 0.6 ± 0.1 | 0.05 ± 0.02 |
| | Processive -> Processive (direction change) | 0.12 ± 0.05 | 0.010 ± 0.005 |
| | Processive -> Immobile | 0.10 ± 0.05 | 0.008 ± 0.004 |
| | Processive -> Diffusive | 0.18 ± 0.06 | 0.014 ± 0.006 |
| | Processive -> Processive (speed change) | 0 | 0 |
| Rich media 30°C + PC190723 | Immobile -> Signal loss | 0.83 ± 0.07 | 0.016 ± 0.002 |
| | Immobile -> Processive | 0.08 ± 0.02 | 0.0015 ± 0.0003 |
| | Immobile -> Diffusive | 0.09 ± 0.02 | 0.0017 ± 0.0004 |
| | Processive -> Signal loss | 0.52 ± 0.05 | 0.021 ± 0.002 |

| | | | |
|---|---|---|---|
| | Processive -> Processive (direction change) | 0.21 ± 0.03 | 0.009 ± 0.001 |
| | Processive -> Immobile | 0.16 ± 0.03 | 0.006 ± 0.001 |
| | Processive -> Diffusive | 0.11 ± 0.02 | 0.0043 ± 0.0009 |
| | Processive -> Processive (speed change) | 0.004 ± 0.004 | 0.0001 ± 0.0001 |
| Rich media 30°C + FtsZ$^{G106S}$ | Immobile -> Signal loss | 0.9 ± 0.1 | 0.033 ± 0.005 |
| | Immobile -> Processive | 0.05 ± 0.02 | 0.0018 ± 0.0008 |
| | Immobile -> Diffusive | 0.008 ± 0.008 | 0.0003 ± 0.0003 |
| | Processive -> Signal loss | 0.6 ± 0.1 | 0.029 ± 0.007 |
| | Processive -> Processive (direction change) | 0.18 ± 0.06 | 0.009 ± 0.003 |
| | Processive -> Immobile | 0.18 ± 0.06 | 0.009 ± 0.003 |
| | Processive -> Diffusive | 0.03 ± 0.02 | 0.001 ± 0.001 |
| | Processive -> Processive (speed change) | 0 | 0 |

**Supplementary Table 3: Fraction of each HT-PBP2B motion state under each experimental condition.** Fractions were calculated by the total amount of time spent in that state relative to the total amount of time spent in any state: $f_{state} = T_{state}/T_{total}$. Standard errors were estimated as $\delta f_{state} = f_{state}\sqrt{\left(\frac{\delta T_{state}}{T_{state}}\right)^2 + \left(\frac{\delta T_{total}}{T_{total}}\right)^2}$, where $\delta T_{state} = \sqrt{N_{state}}$ and $\delta T_{total} = \sqrt{N_{total}}$.

| Condition | Fraction immobile | Fraction processive | Fraction diffusive |
|---|---|---|---|
| Poor media 30°C | 0.567 ± 0.006 | 0.421 ± 0.005 | 0.0116 ± 0.0007 |
| Poor media 37°C | 0.557 ± 0.005 | 0.433 ± 0.004 | 0.0106 ± 0.0005 |
| Rich media 30°C | 0.381 ± 0.004 | 0.589 ± 0.006 | 0.030 ± 0.001 |
| Rich media 37°C | 0.417 ± 0.005 | 0.548 ± 0.006 | 0.035 ± 0.001 |
| Rich media 30°C +penicillin G | 0.955 ± 0.008 | 0.026 ± 0.01 | 0.0195 ± 0.0009 |
| Rich media 30°C +fosfomycin | 0.94 ± 0.01 | 0.040 ± 0.002 | 0.017 ± 0.001 |
| Rich media 30°C +PC190723 | 0.654 ± 0.007 | 0.293 ± 0.004 | 0.053 ± 0.002 |
| Rich media 30°C, FtsZ$^{G106S}$ expression | 0.70 ± 0.02 | 0.290 ± 0.009 | 0.009 ± 0.001 |

**Supplementary Table 4: Primers used in this study.**

| Primer name | Sequence (5'->3') |
|---|---|
| ftsL Fw | ATGAGCAATTTAGCTTACCAACC |

| | |
|---|---|
| spoVD Rev | TCAATCGGCTGCCTCCTTTTC |
| pbp2B Rev | TTAATCAGGATTTTTAAACTTAACCTTGATTACGG |

**Supplementary Table 5: Microscopy acquisition parameters.** N.D. = Not determined.

| Figure number | Microscope configuration | Wavelength (nm) and power density | Exposure time and acquisition interval | Image pixel size |
|---|---|---|---|---|
| 1c | Custom inverted microscope, ring-HiLO illumination | 488: 0.2-0.8 W/cm$^2$<br>561: 7 W/cm$^2$ | 488: 2-5 s exposure, 1 frame<br>561: 500 ms exposure, 1 frame/s | 65 |
| 1d | Custom inverted microscope, ring-HiLO illumination | 488: 0.2-0.8 W/cm$^2$<br>561: 7 W/cm$^2$ | 488: 2-5 s exposure, 1 frame<br>561: 500 ms exposure, 1 frame/s | 65 |
| 1e | Nikon Ti2, HiLO illumination | 488: 7-10 W/cm$^2$<br>561: 11-30 W/cm$^2$ | 488: 5 s exposure, 1 frame<br>561: 500 ms exposure, 1 frame/s | 65 |
| 3a | Custom inverted microscope, ring-HiLO illumination | 488: 0.2-0.8 W/cm$^2$<br>561: 7 W/cm$^2$ | 488: 2-5 s exposure, 1 frame<br>561: 500 ms exposure, 1 frame/s | 65 |
| 3b | Nikon Ti2, HiLO illumination | 488: 7 W/cm$^2$<br>561: 11 W/cm$^2$ | 488: 2-5 s exposure, 1 frame<br>561: 500 ms exposure, 1 frame/s | 65 |
| 4a-d | Custom inverted microscope, ring-HiLO illumination | 488: 0.2-0.8 W/cm$^2$<br>561: 7 W/cm$^2$ | 488: 2-5 s exposure, 1 frame<br>561: 500 ms exposure, 1 frame/s | 65 |
| Supplementary Figure 2a-b | Nikon Ti2, HiLO illumination | 561: 7 W/cm$^2$ | 561: 1 s exposure | 65 |
| Supplementary Figure 2c | Nikon Ti, widefield illumination | 550: N.D. | 550: 200 ms exposure | 65 |
| Supplementary Figure 2d | Nikon Ti2, HiLO illumination | 561: 7 W/cm$^2$ | 561: 1 s exposure | 65 |
| Supplementary Figure 5 | Nikon Ti2, TIRF illumination | 488: 11 W/cm$^2$ | 488: 1 s exposure, continuous | 65 |
| Supplementary Figure 6 | Custom inverted microscope, ring-HiLO illumination | 488: 0.2-0.8 W/cm$^2$<br>561: 7 W/cm$^2$ | 488: 2-5 s exposure, 1 frame<br>561: 500 ms exposure, 1 frame/s | 65 |
| Supplementary Figure 7 | Custom inverted microscope, ring-HiLO illumination | 488: 0.2-0.8 W/cm$^2$<br>561: 7 W/cm$^2$ | 488: 2-5 s exposure, 1 frame<br>561: 500 ms exposure, 1 frame/s | 65 |

| | | | | |
|---|---|---|---|---|
| Supplementary Figure 8 | Custom inverted microscope, ring-HiLO illumination | 488: 0.2-0.8 W/cm$^2$<br>561: 7 W/cm$^2$ | 488: 2-5 s exposure, 1 frame<br>561: 500 ms exposure, 1 frame/s | 65 |
| Supplementary Figure 10 | Custom inverted microscope, ring-HiLO illumination | 488: 0.2-0.8 W/cm$^2$<br>561: 7 W/cm$^2$ | 488: 2-5 s exposure, 1 frame<br>561: 500 ms exposure, 1 frame/s | 65 |
| Supplementary Figure 11 | Custom inverted microscope, ring-HiLO illumination | 488: 0.2-0.8 W/cm$^2$<br>561: 7 W/cm$^2$ | 488: 2-5 s exposure, 1 frame<br>561: 500 ms exposure, 1 frame/s | 65 |
| Supplementary Figure 12 | Nikon Ti2, HiLO illumination | 488: 11 W/cm$^2$<br>561: 7 W/cm$^2$ | 488: 1-3 s exposure, 1 frame<br>561: 500 ms exposure, 1 frame/s | 65 |
| Supplementary Figure 13 | Nikon Ti2, TIRF illumination | 488: 11 W/cm$^2$ | 488: 1 s exposure, continuous | 65 |
| Supplementary Figure 14 | Elyra 7 Lattice SIM2 microscope (Zeiss) | 488: N.D.<br>561: N.D. | 488: 118 ms exposure<br>561: 118 ms exposure | 62 |
| Supplementary Figure 15 | Nikon Ti2, TIRF illumination | 488: 11 W/cm$^2$ | 488: 1 s exposure, continuous | 65 |
| Supplementary Figure 17 | Nikon Ti2, TIRF illumination | 488: 11 W/cm$^2$<br>561: 7 W/cm$^2$ | 488: 1-3 s exposure, 1 frame<br>561: 500 ms exposure, 1 frame/s | 65 |
| Supplementary Figure 18 | Custom inverted microscope, ring-HiLO illumination | 488: 0.2-0.8 W/cm$^2$<br>561: 7 W/cm$^2$ | 488: 2-5 s exposure, 1 frame<br>561: 500 ms exposure, 1 frame/s | 65 |
| Supplementary Figure 19 | Custom inverted microscope, ring-HiLO illumination | 488: 0.2-0.8 W/cm$^2$<br>561: 7 W/cm$^2$ | 488: 2-5 s exposure, 1 frame<br>561: 500 ms exposure, 1 frame/s | 65 |
| Supplementary Figure 20 | Custom inverted microscope, ring-HiLO illumination | 488: 0.2-0.8 W/cm$^2$<br>561: 7 W/cm$^2$ | 488: 2-5 s exposure, 1 frame<br>561: 500 ms exposure, 1 frame/s | 65 |
| Supplementary Figure 21 | Nikon Ti2, HiLO illumination | 561: 7 W/cm$^2$ | 561: 500 ms exposure, 1 frame/s | 65 |

**Supplementary Table 6: Sample sizes.** Number of replicates refers to the number of experiments performed using independently-prepared samples. N.D. = Not determined. *: reproduced here from Whitley et al. 2021[2].

| **Figure number** | **No. data points** | **No. cells** | **No. replicates** |
|---|---|---|---|
| 1f | 1064 track segments | 278 | 4 |
| 1g | 641 track segments | 253 | 4 |

| | | | |
|---|---|---|---|
| 1h | 479 track segments | 154 | 4 |
| 2a, minimal media 30C | 746 track segments | 210 | 3 |
| 2a, minimal media 37C | 1218 track segments | 373 | 3 |
| 2a, rich media 30C | 1064 track segments | 278 | 4 |
| 2a, rich media 37C | 1564 track segments | 461 | 8 |
| 3c | 619 track segments | 252 | 5 |
| 3d | 275 track segments | 77 | 8 |
| Supplementary 2c, Δ*hag* | 185 cells | 185 | 1 |
| Supplementary 2c, 20 µM IPTG | 289 cells | 289 | 1 |
| Supplementary 2c, 50 µM IPTG | 293 cells | 293 | 1 |
| Supplementary 2c, 100 µM IPTG | 254 cells | 254 | 1 |
| Supplementary 2c, 1000 µM IPTG | 262 cells | 262 | 1 |
| Supplementary 2d, 0% xylose | 570 cells | 570 | 1 |
| Supplementary 2d, 0.075% xylose | 592 cells | 592 | 1 |
| Supplementary 2d, 0.375% xylose | 244 cells | 244 | 1 |
| Supplementary 2d, 0.75% xylose | 208 cells | 208 | 1 |
| Supplementary 2d, 1.5% xylose | 114 cells | 114 | 1 |
| Supplementary 4a | N/A | N/A | 3 |
| Supplementary 4b | N/A | N/A | 3 |
| Supplementary 5, SH147 | 85 track segments | N.D. | 3 |
| Supplementary 5, poor media 30°C* | 94 track segments | N.D. | 1 |
| Supplementary 5, poor media 37°C* | 69 track segments | N.D. | 1 |
| Supplementary 5, rich media, 30°C* | 95 track segments | N.D. | 1 |
| Supplementary 5, rich media 37°C* | 87 track segments | N.D. | 1 |
| Supplementary 7 | 206 track segments | 148 | 4 |
| Supplementary 10e | 153 tracks | N.D. | 4 |
| Supplementary 10f | 153 tracks | N.D. | 4 |
| Supplementary 12 | 703 track segments | 193 | 4 |
| Supplementary 13, 21°C | 109 track segments | N.D. | 3 |
| Supplementary 13, 30°C* | 95 track segments | N.D. | 1 |

| | | | |
|---|---|---|---|
| Supplementary 13, 37°C* | 87 track segments | N.D. | 1 |
| Supplementary 14 | 300 cells | 300 | 3 |
| Supplementary 15, WT* | 95 track segments | N.D. | 1 |
| Supplementary 15, G106S | 143 track segments | N.D. | 3 |
| Supplementary 16a, Δhag | 222 cells | 222 | 1 |
| Supplementary 16a, HT-PBP2B GFP-FtsZ Δhag | 318 cells | 318 | 1 |
| Supplementary 16a, FtsZ$^{G106S}$ HT-PBP2B GFP-FtsZ Δhag | 393 cells | 393 | 1 |
| Supplementary 16b | N/A | N/A | 3 |
| Supplementary 17, unperturbed | 189 tracks | N.D. | 3 |
| Supplementary 17, PC190723 | 111 tracks | N.D. | 4 |
| Supplementary 17, G106S | 478 tracks | N.D. | 4 |

**Supplementary Note 1: Calculation of transition rates**

For a state A that can transition to multiple other states B, C, D, etc., where each competing transition is associated with a distinct rate constant $k_B$, $k_C$, $k_D$, etc.:

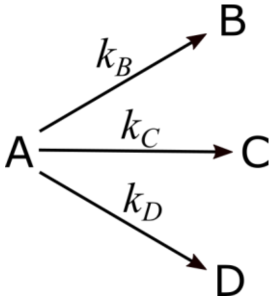

The rate of leaving state A overall is given by a sum of all competing rates:

$$k_A = k_B + k_C + \cdots$$

The rate of depletion of A is given by

$$\frac{dA}{dt} = -k_A A(t)$$

while the rate of accumulation of B is given by

$$\frac{dB}{dt} = k_B A(t)$$

and similarly for the accumulation of other states.

The fraction of state A over time is then given by the first-order ODE as

$$A(t) = k_A e^{-k_A t}$$

and the fraction of state B over time is then given by

$$B(t) = k_B e^{-k_A t}$$

and similarly for the fractions of other states.

If we want to know the total fraction that ended up in state B, we integrate over all time:

$$\int_{t=0}^{\infty} B(t')dt' = \int_{t=0}^{\infty} k_B e^{-k_A t'} dt'$$

yielding

$$B = \frac{k_B}{k_A} = k_B \langle t_A \rangle$$

where $\langle t_A \rangle$ is the mean lifetime of state A.

This means that we can calculate transition rate $k_B$ (and all others) using the fraction of events ending in state B and the measured lifetime of state A:

$$k_B = \frac{B}{\langle t_A \rangle}$$

**Supplementary Note 2: Temperature dependence of FtsZ treadmilling**

We can predict the effect of temperature on the rates of polymerization and depolymerization using the Eyring equation:

$$k = \frac{k_B T}{h} e^{-\frac{\Delta H^\ddagger}{k_B T} + \frac{\Delta S^\ddagger}{k_B}}$$

If we are comparing the rates across two temperatures, we can rearrange this expression to:

$$\frac{k_1}{k_2} = \frac{T_1}{T_2} e^{-\frac{\Delta H^\ddagger}{k_B}\left(\frac{1}{T_1} - \frac{1}{T_2}\right)}$$

along with estimates for $\Delta H^\ddagger$ measured for *E. coli* FtsZ GTPase / depolymerization *in vitro*[7]. Using the published value of $\Delta H^\ddagger$ = 98.4 kJ/mol for depolymerization, we find that an increase from 30°C to 37°C should cause an increase of ~2-fold in this rate, and hence likely in treadmilling speed. Similarly, a decrease from 30°C to 21°C should cause a drop of ~63% in speed.
